# In Silico Investigation of the Role of Local and Global Inflammation-Driven Feedback in Myelopoiesis and Clonal Cell Expansion

**DOI:** 10.1101/2025.08.15.670507

**Authors:** Yusuf Jamilu Umar, Symeon Savvopoulos, Charalampos Pitsalidis, Ioannis Mitroulis, Haralampos Hatzikirou

## Abstract

Chronic inflammation perturbs hematopoietic homeostasis, promoting aberrant myelopoiesis and clonal expansion of mutated stem cells. Here, we develop a mathematical model that integrates both local (bone marrow-intrinsic) and global (systemic/peripheral) inflammation-driven feedback mechanisms to investigate their roles in hematopoietic regulation and disease progression. Our model captures the nonlinear interplay between self-renewal, progenitor proliferation, and inflammatory cues, enabling classification of healthy, myelodysplastic, and leukemic states based on stem cell population dynamics. We show that global inflammatory feedback enhances the resilience of hematopoiesis, while excessive feedback on progenitor cells under chronic inflammation drives instability and clonal dominance. Using sensitivity analysis and parameter space mapping, we identify critical feedback thresholds governing transitions between hematopoietic states and reveal how mutated clones exploit inflammation to outcompete wild-type cells. This systems-level framework offers mechanistic insights into the emergence of myeloid malignancies and provides a computational platform for exploring potential anti-inflammatory therapeutic strategies.

## 1 Introduction

Inflammation is a conserved mechanism that ensures the restoration of homeostasis in response to infection and injury. An important aspect of this mechanism is the response of the hematopoietic system to inflammatory mediators that ensures the demand-adapted replenishment of immune cells. Hematopoietic stem cells (HSC) in bone marrow (BM) can respond to inflammatory stimuli, including pathogen-derived products and inflammatory mediators, resulting in enhanced proliferation and myeloid differentiation [1]. Chronic inflammation, however, leads to loss of HSC fitness, resulting in their depletion ([2, 3, 4]). Aging is considered a chronic inflammatory condition associated with defects in the self-renewal capacity of HSCs, myeloid-priming and increased prevalence of clonal hematopoiesis and myeloid malignancies [5]. Indeed, aging is characterized by a deregulation of the immune system, with enhanced production of inflammatory cytokines that results in a chronic low-dose inflammatory status, which further supports the development and progression of aging-associated disorders, such as cardiovascular disorders and cancer [6].

Emergency granulopoiesis, i.e. the generation of neutrophils in response to inflammation, is a hallmark of acute inflammation [1]. HSC are equipped with a variety of receptors that sense inflammatory stimuli, including Toll-like receptors [7], cytokine receptors [1, 3, 8] and receptors for myeloid lineage growth factors [9, 10], which enables their rapid adaptation to inflammation with enhanced proliferation and myeloid differentiation. However, beyond these direct effects, global feedback mechanisms emerge from systemic inflammatory responses, in which circulating inflammatory signals influence hematopoiesis both in the bone marrow and at peripheral tissues. In the case of chronic inflammation, the exit from dormancy in response to chronic cytokine stimulation, including interferons and interleukin (IL)-1, —forces HSCs into continuous activation, resulting in loss of self-renewal potential, HSC injury, and eventual attrition [3, 8, 11]. Aside from the direct effect of inflammation on HSCs, other cell populations, including stromal cells [12] and granulocytes [13], contribute to global feedback regulation by amplifying inflammatory signaling and further impairing HSC function.

Aging is associated with chronic inflammation, a process termed inflammaging [6, 14], in which systemic inflammatory feedback loops sustain long-term hematopoietic dysregulation, contributing to HSC depletion and functional impairment. Chronic inflammatory stimulation has been linked to the progressive decline in HSC potential, ultimately fostering an environment conducive to clonal hematopoiesis and malignant transformation [5].

### 1.1 Aging inflammation and HSC

The increase in life expectancy in recent decades has resulted in a significant increase in the prevalence of aging-related disorders, which is a major burden on health care systems. Aging-related disorders, including cardiovascular diseases and malignancies, are the leading cause of morbidity and mortality in developed countries, which brings the investigation of aging and aging-related disorders into the focus of interest. Technological development in recent years of next-generation sequencing has revealed the presence in approximately 10% to 15% of individuals over 70 years of age and over 30% at age 85 years of hematopoietic cell clones carrying somatic mutations in Myelodysplastic syndrome (MDS)- and acute myeloid leukemia (AML)-associated genes, including TET2, DNMT3A, ASXL1 and JAK2, with a variant allele frequency (VAF) of ≥ 2%. This condition is termed clonal hematopoiesis of indeterminate potential (CHIP) and it is associated with increased risk for cardiovascular events and a significant increase in the development of MDS or AML [15].

### 1.2 Inflammation and clonal hematopoiesis of indeterminate potential (CHIP)

The increased mortality initially associated with CHIP could not be attributed to the increased risk of progression to MDS or AML [16]. A follow-up study de-identified the clear association between CHIP, coronary heart disease, and early myocardial infarction. Analysis of samples from two prospective cohorts revealed that the risk of coronary heart disease was almost twice as high in people with CHIP compared with the control group. In addition, analysis of retrospective samples of patients with myocardial infarction revealed that patients with CHIP were at four times the risk [17]. Since this initial study, there have been numerous other reports linking CHIP with increased cardiovascular risk and complications of coronary artery disease. For example, the presence of clones positive for ASXL1, TET2 and JAK2 mutations showed a strong statistical association with the occurrence of heart failure independent of the prior presence of coronary artery disease [18].

Progression to CHIP was further associated with modifiable cardiovascular risk factors. In this context, a study was conducted in a sample of 8709 postmenopausal women and correlated the presence of CHIP with lifestyle factors, including body mass index, smoking, physical activity, and diet quality. In this study, CHIP was statistically significantly associated with unhealthy lifestyles, obesity, and smoking. Based on the British Biobank, a cohort of 44,111 individuals was studied and found that the prevalence of CHIP was significantly higher in individuals with unhealthy diets (7.1%), characterized by the consumption of processed foods and high salt intake, compared to individuals with healthy eating habits (5.1%) [19]. These epidemiological findings propose a role for low-grade inflammation associated with the western type of diet and CHIP.

In addition to clinical observations, efforts have been made in experimental models to establish the mechanism explaining clonal hematopoiesis and cardiovascular risk. It has been established that the absence or mutations in the Tet2 gene induce a proinflammatory signature in which IL-1*β* plays an important role, playing a major role in the development of atherosclerosis. These findings provide a link between clonal hematopoiesis, inflammation and cardiovascular disease. On the other hand, studies in inflammatory animal models propose that mutated hematopoietic stem cells are resistant to suppression of their stemness compared to normal clones, resulting in an expansion of mutated clones during inflammation [20, 21]. Taken together, CHIP clones increase the inflammatory burden, whereas inflammation enables such clones to prevail against normal clones, creating a feed-forward loop.

### 1.3 Hematopoiesis and feedbacks

The history of mathematical modeling in hematopoiesis has helped explain blood cell production and disorders. Fokas, Keller, and Clarkson’s mathematical model of granulopoiesis provided a new perspective on chronic myelogenous leukemia (CML) dynamics [22]. Their model explained CML’s paradoxical decrease in proliferation but increased malignant cell production by incorporating physiological parameters, revealing the disease’s pathophysiology. Using feedback-driven dynamics in hematopoietic regulation, Othmer et al. showed how disruptions can cause periodic hematological diseases with blood cell oscillations [23].

Hematological disorders have been better understood through mathematical modeling, as Colijn and Mackey developed comprehensive models for periodic CML and cyclical neutropenia [24, 25]. They simulated disease-specific oscillations and feedback control disruptions using patient and experimental data. Adimy, Crauste, and Ruan developed age-structured differential equation models to describe HSC dynamics and regulation [26, 27]. Their Hopf bifurcation analysis showed how small perturbations could destabilize hematopoiesis and cause disease. Some studies have used lineage specification and mutation-driven dynamics to study the transition from normal hematopoiesis to cancer. Glauche and Peixoto studied how mutations alter hematopoietic differentiation and cause cancer [28, 29]. Silva et al. used multiscale approaches to bridge cellular and spatial scales in the bone marrow [30], while Marciniak-Czochra and Stiehl studied hematopoietic reconstitution and leukemogenesis [31, 32]. Recent advances by Tian and Smith-Miles on gene regulatory networks [33], Klose et al. on aging and progenitor demand [34], and Kumar et al. on leukemia feedback loops demonstrate the evolution and clinical relevance of these models [35].

Hematopoiesis is controlled by complex feedback loops that regulate stem cell self-renewal, differentiation, and proliferation. Hematological malignancies and other disorders result from feedback mechanism disruptions. Rodriguez-Brenes et al. showed how differentiated cell signals inhibit stem cell division and self-renewal, revealing tumorigenesis [36]. Lander et al. used the mammalian olfactory epithelium to study tissue growth feedback loops and optimal differentiation strategies [37]. Kumar et al. studied how feedback configurations affect leukemic cell dynamics using a nonlinear ordinary differential equation model, revealing AML’s aggressiveness [35]. MDS and AML progress due to feedback dysregulation. Walenda et al. found that MDS-initiating HSCs outcompete normal hematopoiesis due to self-renewal [38], while Stiehl et al. found that cytokine-independent proliferation causes early relapse and poor prognosis in AML [39, 40]. Their research shows that patient-specific leukemic cell responses to cytokines affect disease outcomes, and that mathematical modeling can predict therapeutic outcomes.

Normal hematopoiesis and immune function depend on feedback regulation beyond malignancies. Wang et al. showed how inflammation alters hematopoiesis proliferation and promotes disease progression [41]. Inflammation alters hematopoietic lineage commitment to myeloid cells, contributing to trained immunity and myeloid-driven pathologies, according to Chavakis et al. and Matatall et al. [1, 2]. Chronic infections and inflammatory signals like IFN*γ* can suppress bone marrow HSC function, reducing self-renewal and promoting myeloid differentiation, worsening hematopoietic disorders.

Clonal hematopoiesis and leukemia have been studied using mathematical modeling. Johnston et al. examined colorectal cancer’s crypt equilibrium and uncontrolled proliferation to understand hematopoietic transformation [42]. Brunetti et al. noted that clonal hematopoiesis precedes leukemia, emphasizing the need for early detection [43]. Young et al. found that many people have AML-associated mutations, suggesting that clonal expansion above a critical threshold increases leukemia risk [44].

Computational models have revealed stem cell-driven cancers’ regulatory feedback loop escape mechanisms. Rodriguez-Brenes, Komarova, and Wodarz demonstrated how stem cell-driven cancers maintain their regulatory architecture while avoiding growth constraints, emphasizing carcinogenesis’ evolutionary dynamics [45]. These findings help target therapies by identifying cancer cell mechanisms that bypass feedback controls. Roeder and d’Inverno examined mathematical modeling in CML research and inter-disciplinary treatment development [46]. Roeder and Radtke examined stem cell heterogeneity and fate decisions, proving mathematical models can be used in stem cell biology [47]. Roeder et al. approximated agent-based models of hematopoietic stem cell organization using partial differential equations, capturing structural aspects of stem cell age dynamics while reducing computational complexity [48]. Pedersen et al. showed how niche processes affect stem and non-stem cell counts, bone marrow transplantation, clonal competition, and cancer elimination [49].

These studies highlight the importance of feedback regulation in hematopoietic dynamics, clonal evolution, and malignancy progression. Experimental and mathematical methods improve our understanding of hematopoiesis, guiding hematological disorder treatment and precision medicine.

### 1.4 Paper structure

This study develops and characterizes mathematical models of how chronic inflammation regulates hematopoiesis, specifically in the setting of CHIP and its progression to MDS. The disruption of local (bone marrow) and systemic (global) feedback mechanisms that govern HSC lineage patterns under inflammatory circumstances is emphasized. The materials and methods section presents two-tiered hematopoiesis models under inflammatory stress. First, a local feedback model covers bone marrow-intrinsic regulatory mechanisms, whereas a global feedback model accounts for systemic inflammatory effects on HSC and progenitor cell activity. Both models are mathematically formulated and equilibrium states and parameter dependencies are analyzed. Key parameters were adjusted within physiologically realistic ranges to examine their effects on hematopoietic dynamics and illness trajectories without formal sensitivity analysis. Chronic inflammation disrupts hematopoietic control and increases clonal growth, as shown in the results section. We emphasize three important findings: a) Inflammatory stimuli can drive the transition from healthy hematopoiesis to an MDS-like state; b) specific parameter regimes determine transitions between healthy, MDS, and acute myeloid leukemia (AML)-like states; and c) inflammatory conditions enable competition between wild-type and mutant clones, potentially resulting in clonal dominance and malignant progression. Violin plots and parameter space visualizations show how self-renewal, differentiation, and inflammatory feedback affect hematopoietic outcomes. Dysregulated inflammatory feedback promotes clonal hematopoiesis, as discussed in the biological and clinical context. It also shows that competition between two clones—usually a wild-type and an inflammation-favored mutant—can shape hematopoietic cell evolution. Progenitor proliferation rates, niche carrying capacities, and inter-clonal competition coefficients affect malignant clones’ selection advantage. In the conclusion section, mathematical modeling helps explain the complicated interaction between inflammation and clonal dynamics. Future research should combine experimental validation and mutation-specific modeling to increase these frameworks’ prediction power and clinical relevance in hematology.

## 2 Materials and methods

### 2.1 Hematopoiesis model with local feedback (HMLF)

The traditional local feedback hematopoiesis model has three cell populations: stem cells, which can self-renew and differentiate multipotently; progenitor cells, which can self-renew and differentiate, as well; and differentiated cells, which perform tissue functions but cannot proliferate [37, 45]. Also, there are feedback loops, where differentiated cells produce signals that govern stem and progenitor cell behavior. Feedback systems regulate stem cell division, self-renewal, and differentiation to maintain homeostasis. The equations of the HMLF are the following:

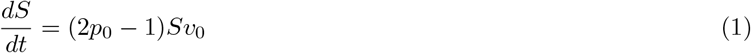

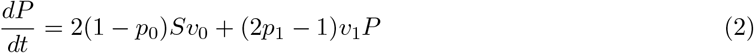

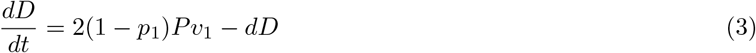

where *S, P*, and *D* denote the populations of stem cells, progenitor cells, and differentiated cells, respectively. The dynamics of these populations are described by eqns 1-3.

The parameter *p*_0_ represents the self-renewal rate of stem cells while *p*_1_ represents the self-renewal rate of progenitor cells. The division rate of the stem cells and progenitor cell populations are denoted by *v*_0_ and *v*_1_ respectively. Finally, the parameter *d* represents the out-flux rate of differentiated cells from the bone marrow compartment into the bloodstream. Here, we interpret *d* as a measure of inflammation affecting the bone marrow environment, where increased values of *d* may be associated with inflammatory states that influence cell turnover. By assuming *d* as inflammation allows us to capture the complex interplay between self-renewal, division, and differentiation, as well as the potential impact of inflammatory conditions on hematopoietic dynamics.

The process is regulated through the secretion of cytokines secreted by various cells within the bone marrow. In this model, the secretion of these proteins depends on the density of the differentiated cells (terminal cells). These proteins regulate the proliferation and differentiation of the stem cells and progenitor cells and are represented by Hill functions. Hill functions are efficient functions used mathematically to capture the sigmoidal relationship between the concentration of a ligand (such as a hormone of substrate) and the response of a biological system behavior.

Mathematically, the feedbacks are given by:

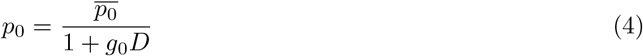

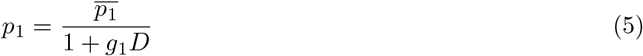

Here, 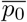 and 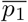 represent the maximal self-renewal fraction of stem and progenitor cells respectively. *g*_0_ and *g*_1_ denote the strength of the negative feedback on the self-renewal rates of stem and progenitor cells respectively.

#### 2.1.1 Parameter estimation of local feedback model

In this subsection, we performed a mathematical analysis of the hematopoietic model at equilibrium, to facilitate the model parametrization. Here, we employ the bone marrow data for the three cellular compartments as reported in [50]. In particular, we relate clinically relevant observables, such as stem cell percentage or progenitor/stem cell ration, cell differentiation parameters. More details are included in sections 2.1 and 2.1.1 of the SI. This analysis enabled us to derive equations for the parameters 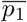 and *v*_1_ as well as to identify constraints that define parameter regimes. These regimes provide critical insight into the progression of the system toward myelodysplastic syndromes (MDS) or acute myeloid leukemia (AML), particularly under perturbations or due to mutations.

Briefly, at steady state, 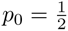, similarly we have

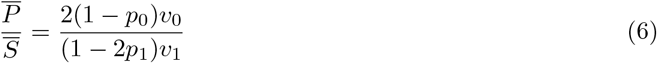

For the steady state of the differentiated cells, we have:

For the steady state of the differentiated cells, we have:

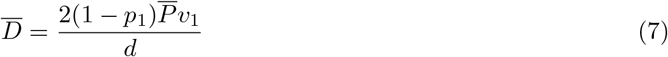

When we consider negative feedback on the model, we can set for simplicity that *g*_0_ = *g*_1_ = *g*, then we have:

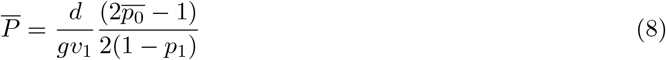

Hence, the ratio of progenitor to stem cells, which is a criterion for the disease stage, becomes:

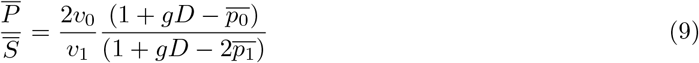

Differentiated cells are responsible for performing specific functions essential for organismal health, such as erythrocytes transporting oxygen, leukocytes defending against infections, and platelets aiding in blood clotting.

Furthermore, through mathematical analysis of the model, assuming *g*_0_ ≠ *g*_1_, we further derived some estimates for the values of the parameters *v*_1_ and *p*_1_. Furthermore, we derived constraints that enabled the identification of parameter ranges associated with the progression to myelodysplastic syndrome and acute myeloid leukemia.

From eqns (1) and (2), assuming 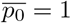, we have at steady state:

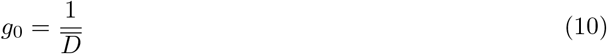

If we assume that 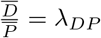 represents the ratio of differentiated cells to that of progenitors at steady state and 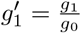, we now have:

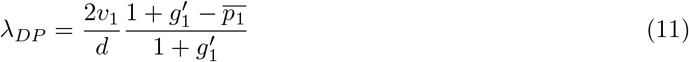

resulting in:

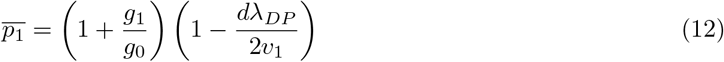

From eq. (12), we can see that since *g*_0_ and *g*_1_ are positive and *p*_1_ ≥ 0, then we derived the constraint

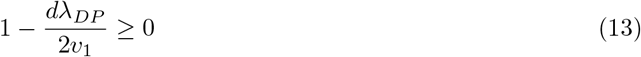

which implies:

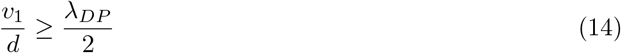

In a similar manner, under the assumption of steady-state conditions, where the ratio of progenitor cells to stem cells is denoted by *λ*_*PS*_, the parameter *v*_1_ is found to be directly proportional to the outflux of differentiated cells. This relationship is expressed mathematically as follows:

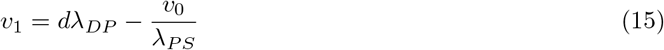

### 2.2 Hematopoiesis model with global feedback (HMGF)

Feedback control is crucial for preventing overgrowth, preserving the stem cell pool, and responding to physiological demand, such as injury or stress, by limiting cell proliferation and allowing the system to dynamically adjust production. The traditional hematopoiesis model overlooks the impact of external systemic factors like inflammation on hematopoietic dynamics [5, 41]. Inflammation, particularly under stress, disrupts this balance, leading to depletion of stem cells, clonal hematopoiesis, and progression to malignancies. The newly extended model integrates global inflammation as a dynamic regulatory factor, significantly enhancing its robustness in predicting hematopoietic dynamics under inflammatory conditions. By accounting for systemic inflammatory effects, the model ensures stability across varying inflammatory states, accurately capturing disease progression, and potential therapeutic intervention points. This framework provides a more resilient and comprehensive tool for studying hematopoiesis, particularly in pathological scenarios driven by chronic inflammation

The model is as follows:

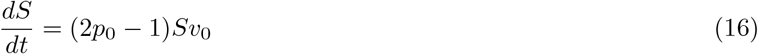

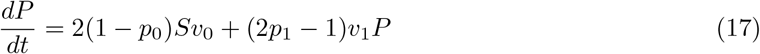

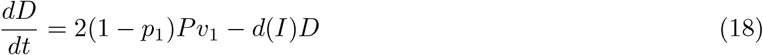

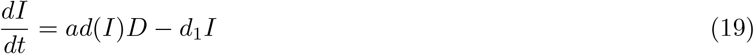

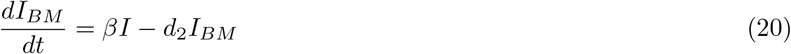

Where *S, P* and *D* are as before, representing the respective cell populations. Here myeloid differentiated cells are typically innate immune cells such as macrophages, neutrophils, etc. Inflammation in the bloodstream periphery is represented by *I* while *I*_*BM*_ inflammation is in the bone marrow compartment. Parameters *a, d*_1_, and *d*_2_ are quantifying the dynamics of inflammation within both the periphery and bone marrow.

The parameter *a* represents the rate at which differentiated cells exit in the bloodstream and subsequently impact peripheral inflammation. The parameter *d*_1_ represents the transport of pro-inflammatory factors, such as cytokines, from the bloodstream into the bone marrow. The latter proinflammatory factors increase the bone marrow inflammation by a rate *β*. Lastly, *d*_2_ represents the rate of decay of inflammation in the bone marrow compartment.

Interestingly, the *d* parameter in eq. (18) is a function of systemic/peripheral inflammation:

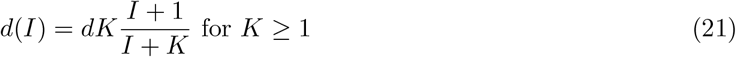

The *d*(*I*) represents a quasi-steady state behavior of the outflux of mature cells from bone marrow. This term models the fact that increased periphery inflammation, due to pathogen attack, requires enhanced immune cell recruitment. In response to the latter, the bone marrow releases more differentiated immune cells. Therefore, we model this *d*(*I*) which is monotonous in *I* with a saturating behavior. The parameter *K* in the expression is a tuning parameter that acts as an inflammation threshold and has *inflammation units*. We can see that when *I* = *K, d*(*I*) reaches approximately half of its asymptotic limit *dK*, e.g. for *I ≫ K*. This saturating behavior is commonly used in the dynamics of cellular signaling, where response variables or intermediate populations quickly settle into a stable relationship with their driving variables. Therefore, the parameter *d*(*I*) ranges from the value *d* to *dK* for *I* ∈ [0, +∞). The latter implies that in the absence of systemic inflammation (*I* = 0), the differentiated cell release rate is *d* and for high inflammation reaches *dK*.

After the above model modifications, the negative feedbacks on stem cells and progenitor cells selfrenewal rates are represented mathematically as:

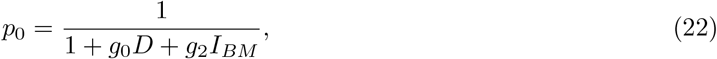

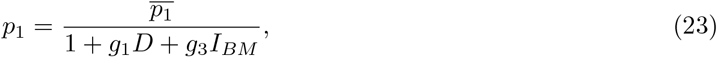

where *g*_0_ signifies the sensitivity level of the negative feedback on stem cell self-renewal influenced by the density of mature cells, *g*_1_ indicates the sensitivity of the negative feedback affecting progenitor cell self-renewal owing to the density of mature cells, *g*_2_ denotes the sensitivity level of stem cell self-renewal impacted by bone marrow inflammation, and *g*_3_ signifies the sensitivity level of progenitor cell self-renewal as a result of bone marrow inflammation.

This extended model of hematopoiesis, which integrates feedback loops dependent on inflammation, offers a comprehensive framework for examining both normal and malignant clonal dynamics under identical inflammatory conditions. This methodology, in contrast to traditional models, elucidates the distinct responses between normal and clonal hematopoietic populations to inflammation. The model illustrates emergency myelopoiesis in normal hematopoiesis, where inflammation influences immune homeostasis. In malignant contexts, it underscores how inflammation alters healthy and mutated clones, promoting self-renewal and advancing disease progression. This dual functionality facilitates the exploration of the impact of inflammation on hematopoiesis and the evaluation of therapeutic strategies.

#### 2.2.1 HMGF model reduction under quasi-steady state assumption

The above model extension introduces a number of new parameters that is difficult to measure in a clinical setting. To address this, we attempt a model reduction by assuming the multiscale nature of our system. In particular, we will use the quasi-steady state approximation to reduce model dimensions and parameters. More details on its analysis can be found in Sections 2.2 and 2.2.1 of the SI. Briefly, at steady state, from eqns (19) and (20), we have:

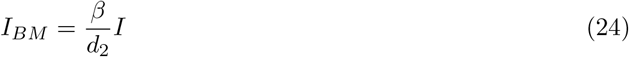

and

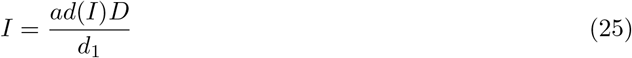

Both eqns (24) and (25) represent quasi-steady-state relations, with soluble factors, such as cytokines, exhibiting higher relaxation characteristic times than their cellular counterparts. Another important observation is that inflammation *I* is proportional to the myeloid differentiated cells.

Assuming that the outflux of the terminal cells is governed by a monotonically increasing function (eq. (21)).

Let 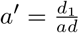 we have:

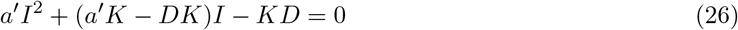

Solving the quadratic equation, we obtain:

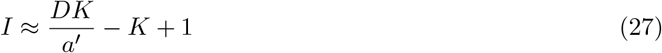

From eq. (27), we see that for large *D*, the dominant term 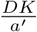 scales linearly with *K*. Thus the term that describes the exit of differentiated cell to the periphery reads

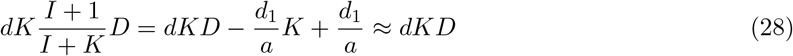

Since *D* order of magnitude is in 10^4^, we take into account only the leading term. Hence, the system of eqns (16)-(20) can be modified to the new system:

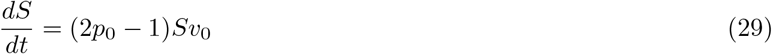

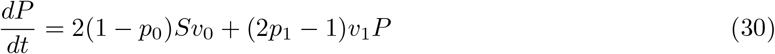

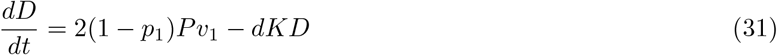

Here, the model complexity is reduced from five to three ODEs with the *K* parameter playing an important role and quantifying the inflammation degree in the periphery. After the aforementioned model reduction, the corresponding negative feedbacks from eqns (22) and (23) are modified as

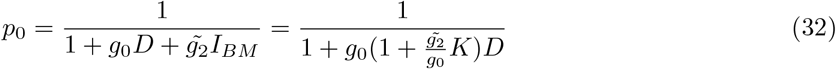

and

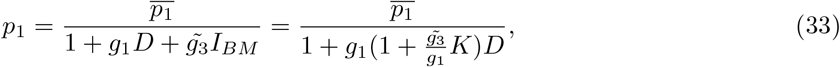

where 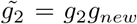 and 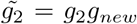. For complete details of the analysis, refer to section 3 of the Supplementary Information.

### 2.3 Healthy and malignant clone competition model

To gain a comprehensive understanding of the interactions and competitive behaviors between two distinct cell populations, a healthy cell population and a leukemic cell population, we employed a competition-based modeling approach. This framework was specifically designed to elucidate how the growth and proliferation of one population influence the dynamics of the other. By integrating competition terms into the kinetic equations, the model captures the inhibitory effects exerted by each population on the other, thereby simulating the complex biological interactions that arise in such systems. This approach allows for the exploration of critical parameters, such as growth rates, carrying capacities, and interpopulation competition coefficients, providing valuable insights into the dynamic balance between healthy and cancerous cell populations. Furthermore, the model serves as a basis for understanding the resource limitations and competitive pressures that govern their coexistence or dominance under various conditions.

The model is similar to the one presented in Section 2.2, which addresses global feedback with chronic inflammation, and its equations are as follows:

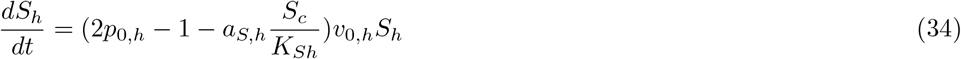

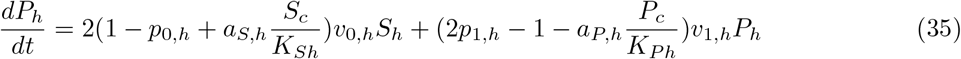

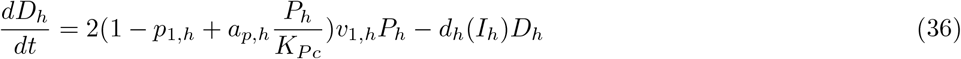

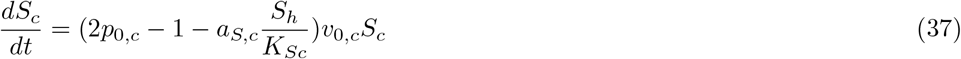

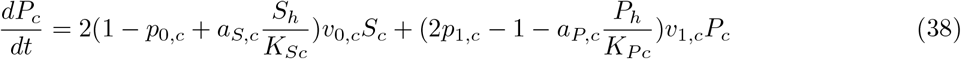

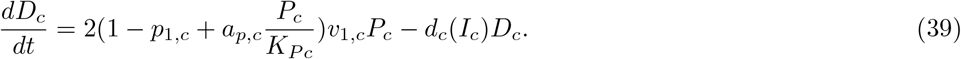

The subscripts *h* and *c* represent the healthy and cancerous populations, respectively. The parameters *a*_*S*_ and *a*_*P*_ denote the competition coefficients for the stem cell and progenitor populations, while *K*_*S*_ and *K*_*P*_ represent their respective carrying capacities.

### 2.4 Classification of disease states in hematopoiesis

To classify the different disease states in hematopoiesis, we ran the model with static inflammation using parameter sets generated from the Sobol function and a grid search method. These parameter sets satisfied the conditions outlined in section 2.1.1. Based on these simulations, the following criteria were established for the percentage of stem cells:

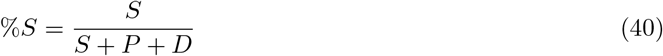

In healthy states, the percentage of stem cells ranges between 1% and 3%, indicating a balanced and functional hematopoietic system. For MDS, the percentage increases to a range of 3% to 20%, reflecting abnormal differentiation and proliferation patterns. In AML, the percentage exceeds 20%, signifying a pathological condition with impaired hematopoiesis and dominance of immature cells. These thresholds provide a clear framework for distinguishing between healthy and diseased states based on the percentage of stem cell values.

### 2.5 Statistical analysis

Descriptive data were analyzed using Microsoft Excel (Office 365, Microsoft) and are provided as mean *±* standard deviation 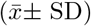. Statistics analysis was conducted using Python with the Pingouin open-source statistics program [51]. A Kruskal-Wallis test was performed to assess significant differences in biomarker levels among the groups [52]. A Mann-Whitney post hoc test was employed to compare differences between any two groups [53]. Significant data were characterized by a p-value of less than 0.05.

### 2.6 Sensitivity analysis

Global sensitivity analysis is crucial for understanding the influence of input parameters on the outcomes of complex models, particularly in nonlinear systems. The random sampling-high dimensional mathematical representation (RS-HDMR) method offers a comprehensive framework for this purpose, enabling an in-depth exploration of parameter sensitivity [54]. More details regarding the mathematics of RS-HDMR analysis and how it works are included in Section 3 of the SI.

RS-HDMR evaluates the sensitivity of parameters across their entire range, unlike local sensitivity analysis, which focuses on small deviations around a fixed point. This global approach ensures that the full scope of parameter interactions and nonlinearities is considered, making it particularly suited for models with complex dynamics.

A key feature of RS-HDMR is its ability to decompose model outputs into hierarchical components. These include the first order effects which shows the direct influence of individual parameters, Second order effects which estimate the interactions between pairs of parameters and the higher-order effects which capture the complex interactions among multiple parameters

This decomposition is achieved through an analysis of variance (ANOVA)-like approach, which allows for a detailed breakdown of how inputs contribute to output variability [54]. More details are included in Section 4 of supplementary material.

RS-HDMR calculates partial variances (*D*_*i*_, *D*_*ij*_) and sensitivity indices (*S*_*i*_, *S*_*ij*_) to attribute output variability to specific parameters or their interactions. These metrics are invaluable for identifying the most critical parameters that require precise measurement or control, optimizing resource allocation, and improving model reliability.

The versatility of RS-HDMR makes it indispensable in diverse fields such as environmental modeling, engineering, and biology [55, 56, 57, 58, 59]. Its ability to handle high-dimensional inputs and quantify parameter importance enhances the accuracy and robustness of models, leading to more informed decision-making processes.

Nominal values and initial conditions used for the implementation of the models can be found in Section 4 of the SI.

## 3 Results

This section analyzes hematopoietic dynamics under chronic inflammation and clonal proliferation using mathematical model and computational simulation results. The results have three main subsections for study aspects. Section 3.1 explains how inflammation causes MDS from healthy hematopoiesis. Section 3.2 introduces complex inflammation effects to reveal key thresholds and model parameter interactions in the parameter space driving healthy, MDS, and AML state transitions. Section 3.3 presents parameter space analysis of how feedback strengths affect transitions between MDS and AML, highlighting that excessive feedback on pluripotent cells may accelerate disease progression. Section 3.4 uses sensitivity analysis to identify critical parameters in healthy and malignant cell population competition. These subsections explain how inflammation and clonal competition affect hematopoietic system dynamics and cancer progression.

### 3.1 Inflammation can lead to myeloid pathologies in the absence of global feedback

The traditional local feedback hematopoiesis model comprises three cellular populations: stem cells, capable of self-renewal and multipotent differentiation; progenitor cells, which also possess self-renewal and differentiation abilities; and differentiated cells, which execute tissue functions but lack proliferative capacity (Figure 1) [37, 45].

**Figure 1.**
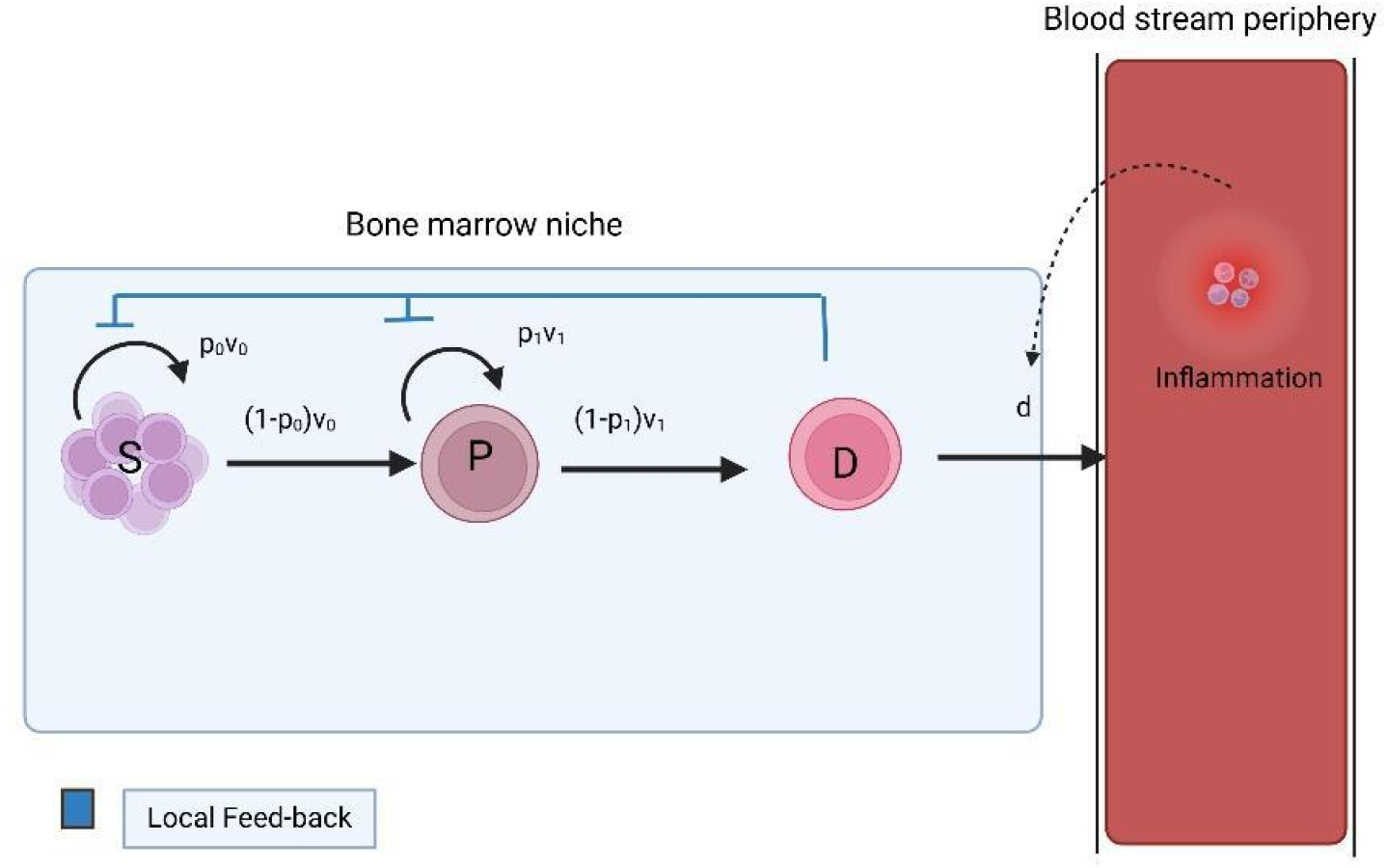
Schematic diagram showing the hematopoietic model with local feedback in the bone marrow.

Initially, we employ the hematopoietic model with local feedback on the stem cell self-renewal, as a control for assessing the myelopoiesis robustness under parameter perturbations. In particular, we are interested in which parameter ranges to observe Healthy, MDS, and AML. This initial modeling step was critical for establishing the validity of parameter correlations with the studied states.

All model parameters were constrained within predefined ranges, with the exception of the parameter *g*_0_, which was held constant as defined by eq. (10). The parameter *g*_1_ was allowed to vary between 0.1 *×* 10^−5^ and 0.9 *×* 10^−5^, while *v*_0_, *v*_1_, and *d*, which are probabilities, were each varied within the range of 0.1 to 0.9. Nominal values of the parameters are in Table 1 of SI. These probability ranges were selected to reflect biologically plausible values and to enable a comprehensive exploration of the parameter space [50].

**Table 1.** Classification of disease state based on stem cell percentage.

| Disease State | Percentage of stem cells(%) |
| --- | --- |
| Normal hematopoiesis | 1 – 3 |
| Myelodysplastic syndrome (MDS) | 3 – 20 |
| Acute Myeloid Leukemia | > 20 |

To identify the parameter sets that aligned with the desired hematopoietic states, it was necessary to satisfy the constraints and equalities outlined in eqns (10) - (15). These mathematical conditions ensured that the selected parameters adhered to the dynamic relationships inherent to the hematopoietic system. The findings of this analysis are illustrated in Figures (2) and (3), which display violin plots representing the distribution of parameter values, cell counts, and the percentage of stem cells among various groups. The violin plots are produced by methodically altering the ranges of the key parameters using Sobol sampling, enabling the examination of how various parameter combinations relate to different hematopoietic states, including healthy, MDS, and AML. This method captures the variability in disease progression and offers insights into the transitions between these states. Moreover, Table 2 in Section 5 of SI presents comprehensive statistical summaries for each parameter, thereby reinforcing the differences and trends identified in the model.

**Figure 2.**
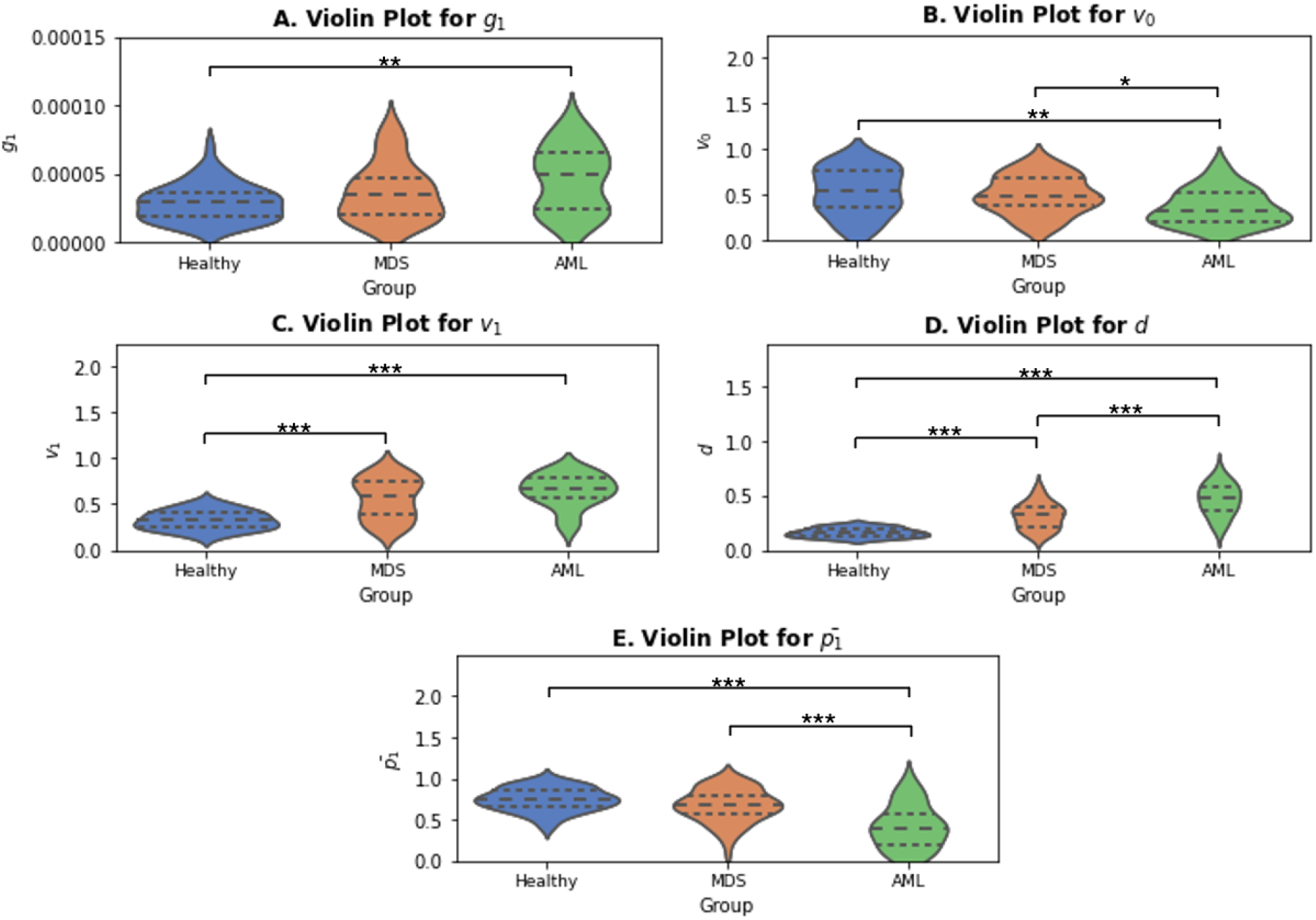
Violin plots with Bonferroni corrected Mann-Whitney statistical analysis of the parameters A) *g*_1_, B) *v*_0_, C) *v*_1_, D) *d*, and E) 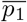 for different disease states (*-*p <* 0.05; ** −*p <* 0.01; ***-*p <* 0.001). Kruskal Wallis values were significant in all parameters.

**Figure 3.**
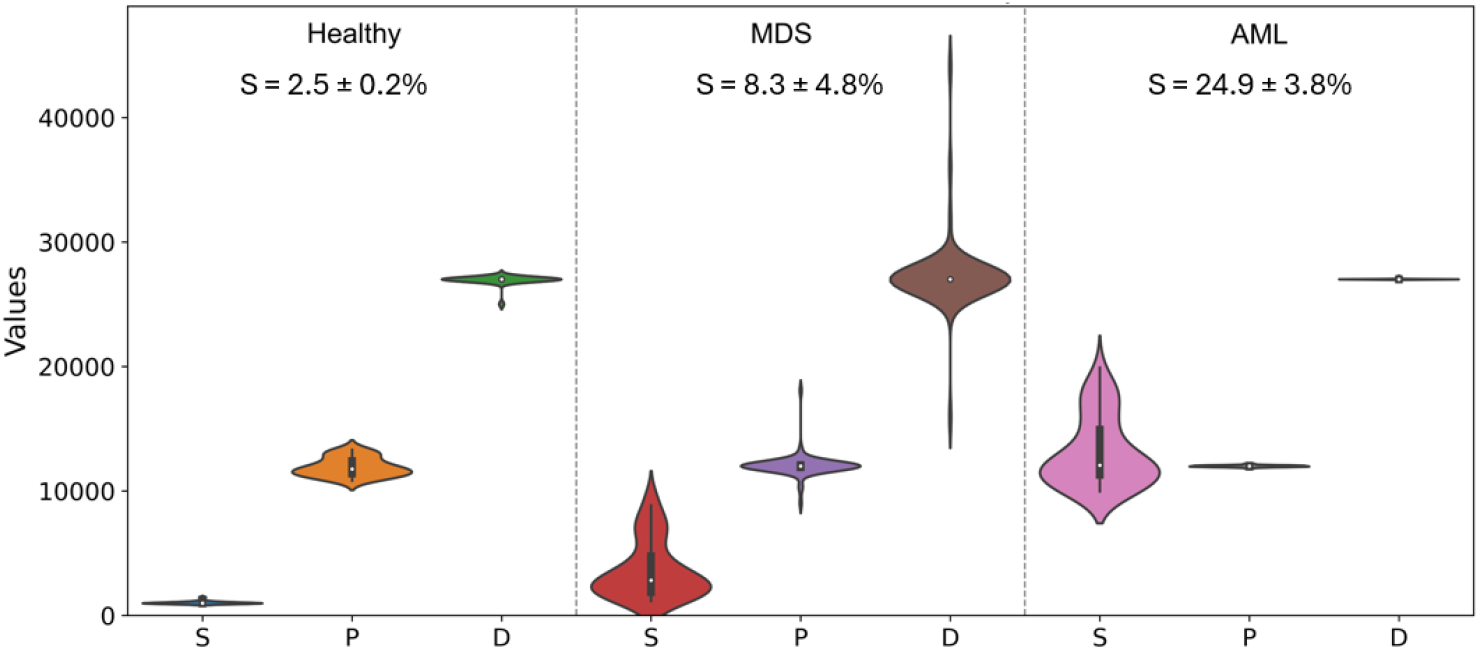
Violin plots of the number of S, P, and D cells in each disease stage.

Our analysis reveals key differences among the healthy, MDS, and AML groups across multiple biological parameters, each reflecting distinct aspects of the system’s state. The violin plots (Fig. (2)) illustrate the distribution of these parameters, while the summary statistics provide a quantitative understanding of the observed variations. Most parameters display distinct patterns across the groups, pointing to potential changes in biological functions or systemic responses associated with disease progression.

Specifically, *g*_1_ demonstrates a slight upward shift in both mean and variability from healthy to MDS and AML classes, suggesting alterations in its regulatory role. Similarly, the proliferation rates of stem cells (*v*_0_) and progenitor cells (*v*_1_) exhibit divergent trends: *v*_0_ decreases significantly in AML, whereas *v*_1_ shows a marked increase, potentially reflecting different pathways or mutations activated in advanced disease. Additionally, the 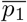 decreases notably in the AML group, implying disruptions in local feedback mechanisms.

Among all parameters, *d* stands out as a critical indicator of systemic inflammation, since it models the mobilization rate of differentiated myeloid cells (e.g., neutrophils) towards the periphery. The demands for more myeloid cells increase during systemic inflammation. The violin plots illustrate a pronounced progression in *d* values, with its distribution shifting markedly upward from the healthy to MDS and then to AML groups. This pattern is corroborated by the mean values (see SI Table 2 of Section 5), where d increases from 0.17 in healthy to 0.32 in MDS and reaches 0.46 in AML cases. The increasing standard deviation, 0.04, 0.12, and 0.14, respectively (see SI Table 2 of Section 5), suggests growing heterogeneity in the inflammatory response, likely reflecting the complexity and variability of disease states.

### 3.2 Inflammation feedback on HSC self-renewal enhances the robustness of hematopoietic homeostasis

The traditional hematopoiesis model neglects the influence of external systemic factors, such as inflammation, on hematopoietic dynamics. Inflammation, especially during periods of stress, disturbs this equilibrium, resulting in stem cell depletion, clonal hematopoiesis, and increased risk for malignancies. The newly expanded model incorporates global inflammation as a dynamic regulatory element, markedly improving its reliability in forecasting hematopoietic dynamics in inflammatory contexts (Figure (4)).

**Figure 4.**
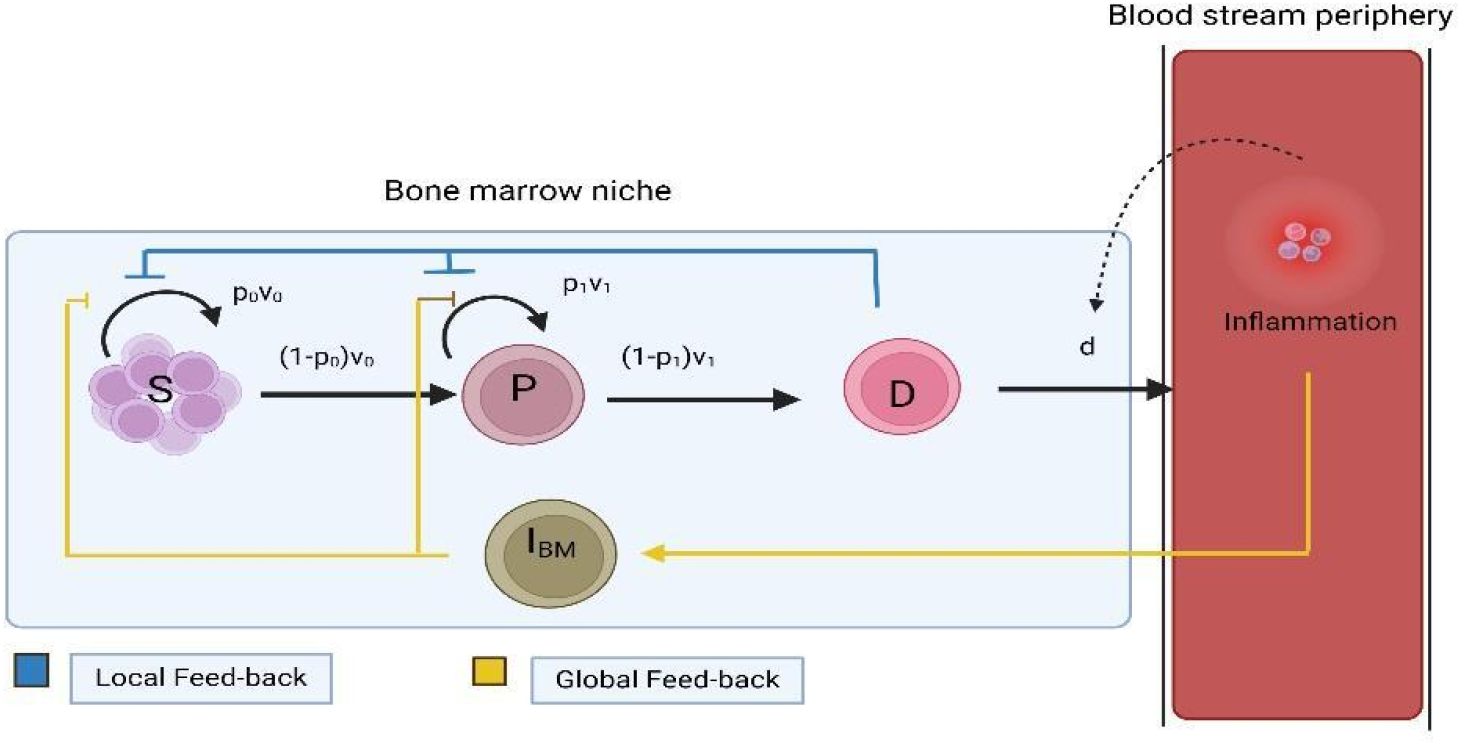
Schematic diagram showing the hematopoietic model with global feedback inflammation in the bone marrow.

In the HMGF, systemic inflammation is explicitly incorporated through the parameter *K*, which is introduced into the mobilization rate *d*(*I*) (see Materials and Methods for details). This mobilization rate governs the dynamics of cell recruitment or movement and depends on the presence and intensity of inflammatory signals in the bloodstream. Notably, *K* serves as an indicator of the onset of inflammation, marking the initial activation or triggering of inflammatory pathways.

The parameter *K* plays a pivotal role in initiating the inflammatory response by influencing *d*(*I*), thereby linking early-stage inflammatory signals with subsequent migration processes. Its inclusion captures how the emergence of inflammation alters baseline conditions, effectively triggering the dynamic interplay between inflammation and mobilization. Under the quasi-steady state assumption, described in

Section 2.2.1, one can show that the degree of inflammation is proportional to the parameter *K*. There on, we will use the parameter *K* as a proxy for the degree of inflammation in the periphery.

Then, we focus on the effect of stem cell self-renewal feedback (global) on hematopoietic pathologies under different degrees of systemic inflammation. To investigate the progression to hematopoietic malignancies, we assumed that there is no inflammation feedback on the progenitor cells, i.e., the parameter *g*_3_ was fixed at zero (*g*_3_ = 0). Keeping all other parameters to healthy values (as identified in the previous section), we varied the inflammation degree *K* and the associated self-renewal feedback (global) strength 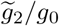 and recorded the resulting stem cell percentage to assess the state of the hematopoietic process.

Figure (5) illustrates the relationship between the inflammation and global feedback, along with their influence on the healthy and MDS system’s states. When the global feedback values are high, inflammation cannot deteriorate the healthy state. However, for lower feedback strengths, there is an associated threshold of inflammation intensity that leads to MDS. This exhibits the protective role of global feedback by enhancing the hematopoiesis robustness under inflammation perturbations.

**Figure 5.**
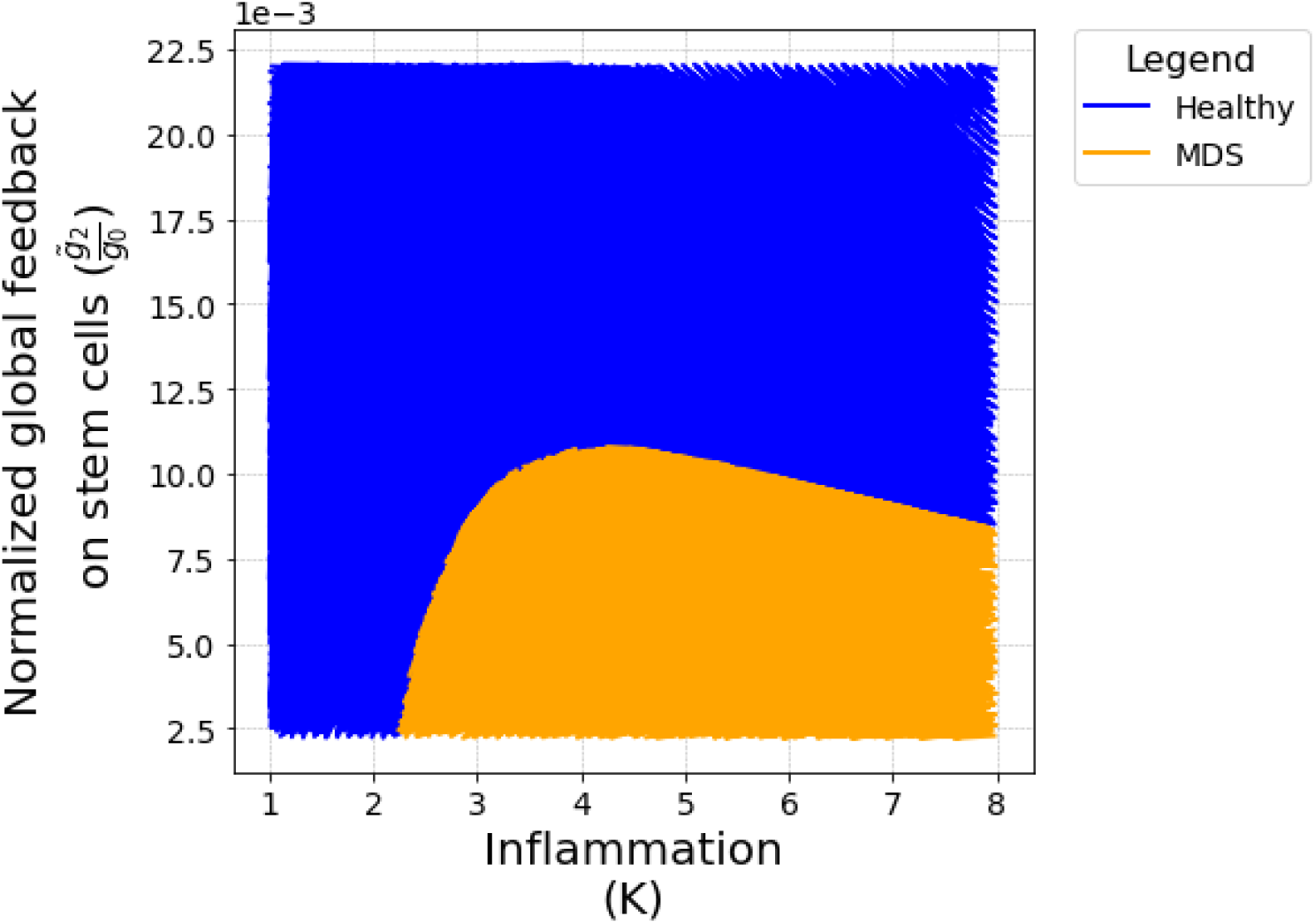
Parameter space analysis of inflammation (*K*) and the normalized global feedback on SC 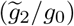 in determining healthy and MDS states.

Figure (6) illustrates the distribution of cell populations in the healthy and MDS states shown in Figure 5. Global feedback controls efficiently differentiated cells and stem cells population, exhibiting a tight variance under inflammation variations. On the other hand, progenitor cell distribution shows a high range both for healthy or MDS states for the associated 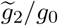 and *K* ranges. In the healthy state, progenitor cells exhibit a long tail, extending to nearly 100000, indicating that progenitor cells act as a buffer when inflammatory conditions change. This is the trade-off in controlling tightly stem cell and differentiated myeloid cell numbers.

**Figure 6.**
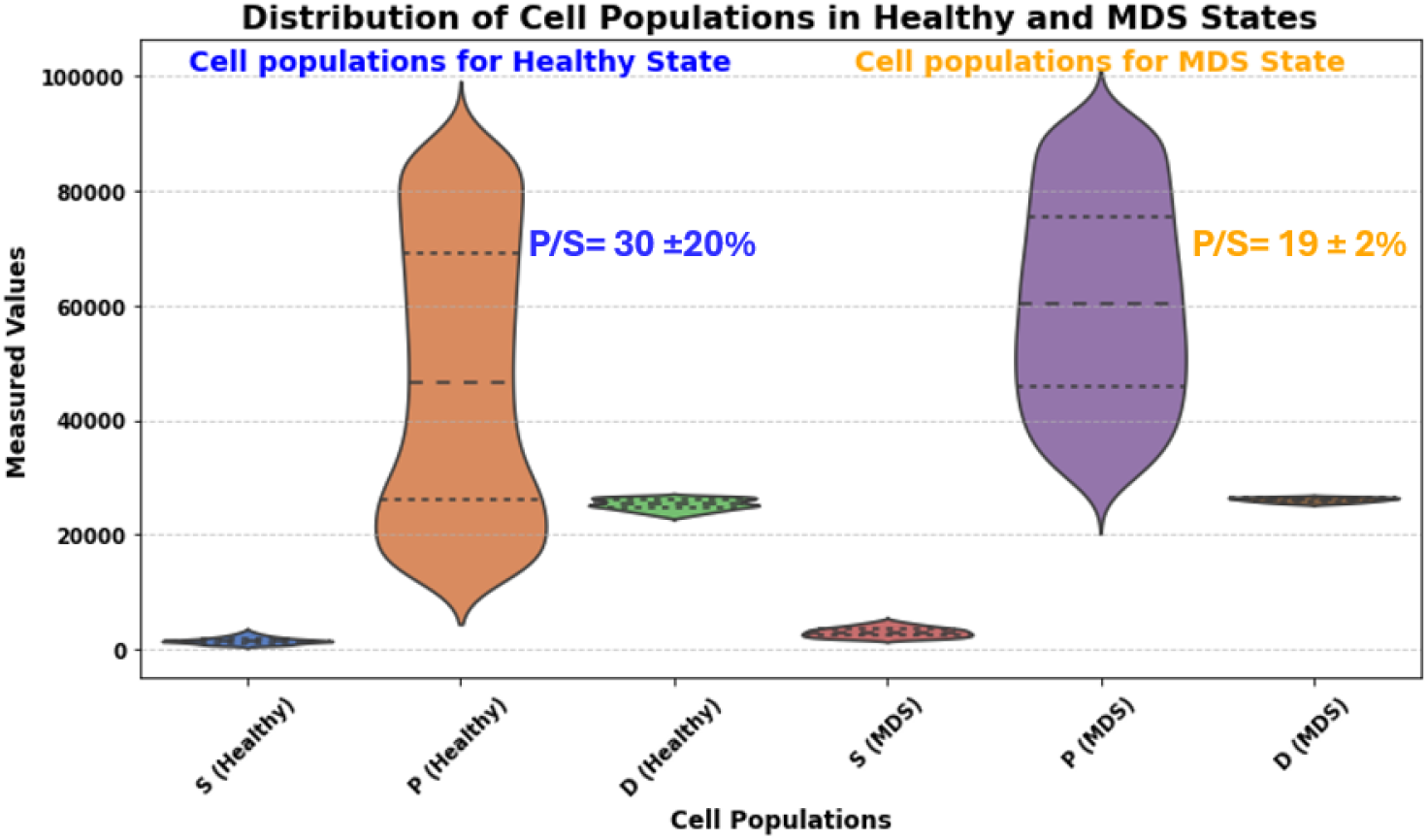
Distribution of cell populations in healthy and MDS states under inflammation (*K >* 1)

Importantly, under inflammatory conditions (*K >* 1), the progenitor-to-stem cell (P/S) ratio changed significantly. In a healthy state, the P/S ratio is around 30 *±* 20%, indicating a significant progenitor cell expansion compared to stem cells. In contrast, this ratio drops to 19 *±* 2% in the MDS state.

This reduction indicates that excess inflammation disrupts the balance between stem and progenitor cells, leading to a relative suppression of stem cell activity. The effect of inflammation is directly visible in the narrowed and lowered distribution of stem cells in MDS, alongside a maintained or slightly elevated progenitor population. This P/S shift shows how inflammation biases hematopoiesis toward progenitor cell expansion at the expense of stem cell maintenance, even though differentiated cell distribution remains relatively unchanged across states. These changes in progenitor and stem cell proportions cause MDS-related hematopoiesis dysfunction.

### 3.3 Increased inflammation feedback on progenitor cells can deteriorate myelopoiesis homeostasis

In this section, we study the effect of global feedback on progenitor cell self-renewal. In particular, we vary the stem cell feedback parameter 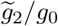 around its healthy values, while we vary the progenitor cell feedback 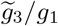 for four orders of magnitude (10^−4^, 10 ^0^). In turn, we vary the inflammation parameter *K*, ranging from 1 to 5, and we assess the hematopoiesis states when stem and progenitor cell feedback strengths are modulated. The three subplots of Figure (7) show three distinct regions corresponding to healthy (blue), MDS (orange), and AML (red) states.

**Figure 7.**
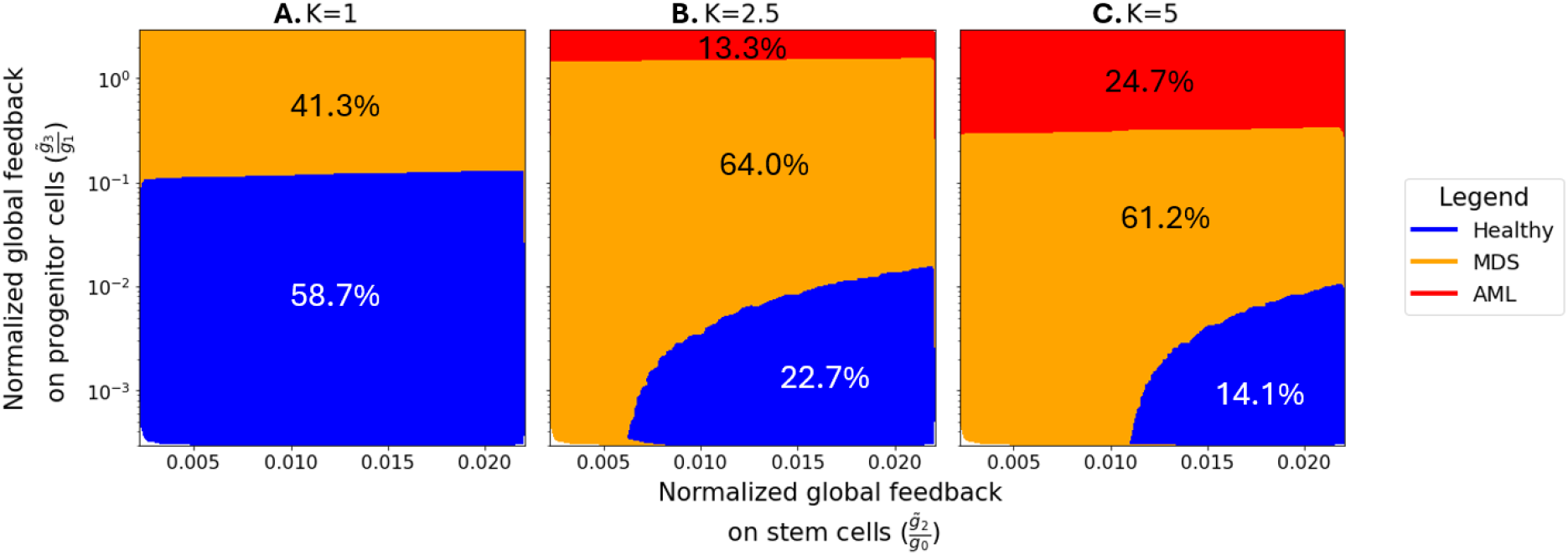
Parameter space analysis of 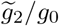 and 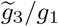 in determining healthy, MDS and AML transitions for different values of the inflammation parameter *K*: A) *K* = 1, B) *K* = 2.5, and C) *K* = 5.

Figure (7)A shows that for no inflammation (*K* = 1), we can obtain 58.7% healthy myelopoiesis and with 41.3% MDS. The absence of AML in this case indicates that very low inflammation does not favor the emergence of malignancies even when progenitor feedback attains high values.

For intermediate inflammation values *K* = 2.5 (Figure (7)B), the healthy region decreases to 22.7% and MDS increases to 64.0%, indicating a higher susceptibility to dysregulated hematopoiesis when global feedbacks are perturbed. The system approaches critical instability when the progenitor feedback 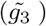 is much higher than its stem cell counterpart 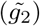, where small but notable AML region (13.3%) emerges. The distinction between healthy and MDS becomes clearer, highlighting K’s role in promoting preleukemia. For high inflammation *K* = 5 (Figure (7)C), the AML region increases to 24.7%, indicating a higher risk of leukemia. MDS dominates 61.2% of the parameter space, while the healthy region shrinks 14.1%. Deregulated self-renewal feedback puts the hematopoiesis process at high pathological risk, especially under high chronic inflammation.

Figure (8) presents violin plots depicting the distribution of cell populations across healthy, MDS, and AML states when *K* = 5. In the healthy state, progenitor cells exhibit a balanced distribution, with values peaking around 20000 to 30000, indicating a stable and regulated population. Stem cells show a moderate distribution, with measured values primarily under 1500, reflecting their supportive role in normal hematopoiesis. Differentiated cells display the smallest distribution, with values tightly clustered below 10000, suggesting a controlled presence in healthy conditions.

**Figure 8.**
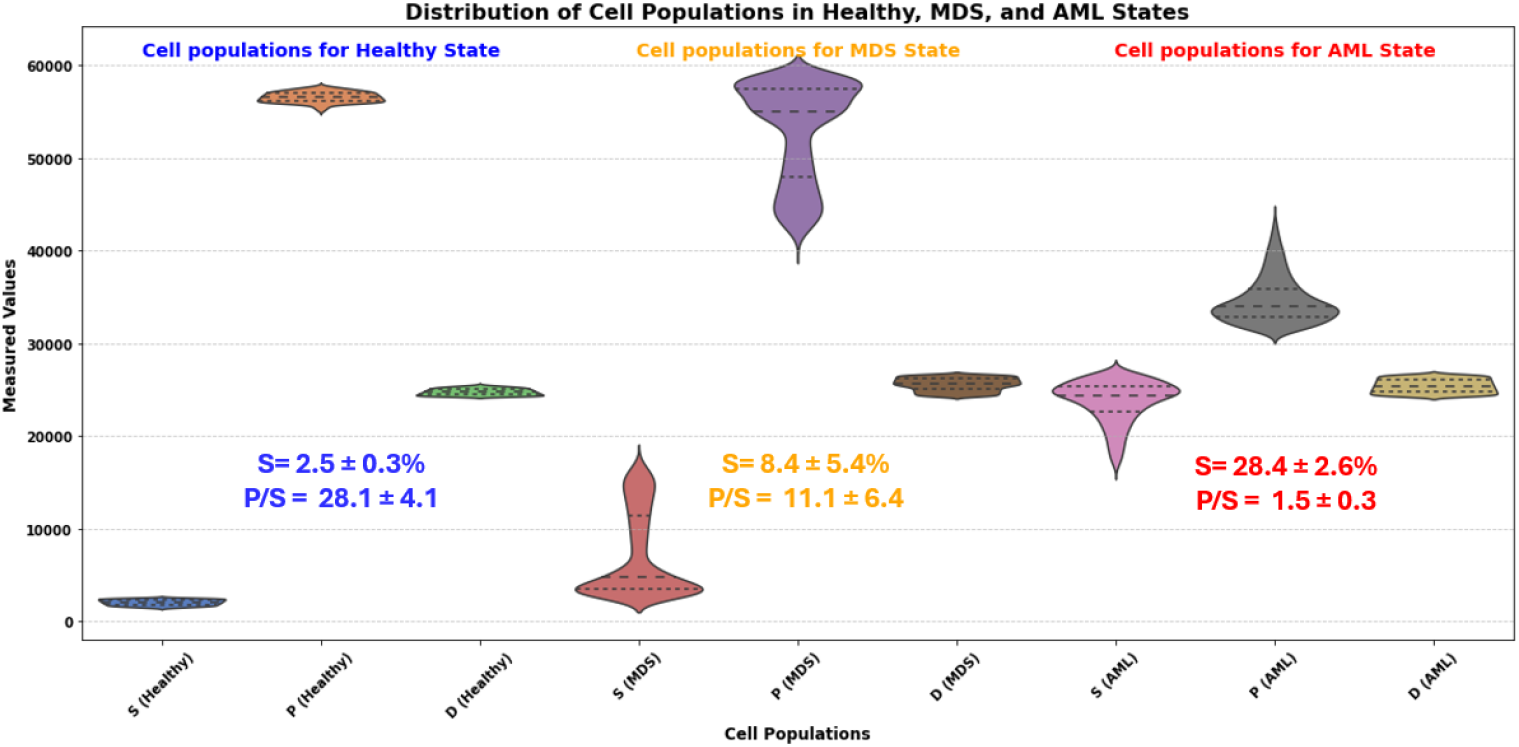
Distribution of cell population in Healthy, MDS and AML states.

Transitioning to the MDS state, the stem cell distribution is mainly concentrated around the healthy values but exhibits a long tail towards higher values. Similarly, progenitor cell distribution is also clustered around the healthy mean value while showing a long tail down to 40000 cells. Interestingly, the differentiated cells stay tightly regulated around their healthy counterparts.

In the AML state, progenitor cells dominate with an even broader distribution, extending up to 60000, reflecting further pathological expansion characteristic of AML progression. Stem cells in AML display an increased distribution compared to MDS, with measured values peaking between 20000 and 30000, indicating significant changes in their population dynamics. Differentiated cells remain suppressed in AML, showing a narrow and low distribution, consistent with observations in healthy and MDS states.

### 3.4 Clones with impaired self-renewal feedback outcompete healthy ones resulting in MDS development

The competition-based model captures the dynamic interactions between healthy and leukemic cell populations by incorporating growth rates, carrying capacities, and competition terms; these interactions are demonstrated in Figure (9).

**Figure 9.**
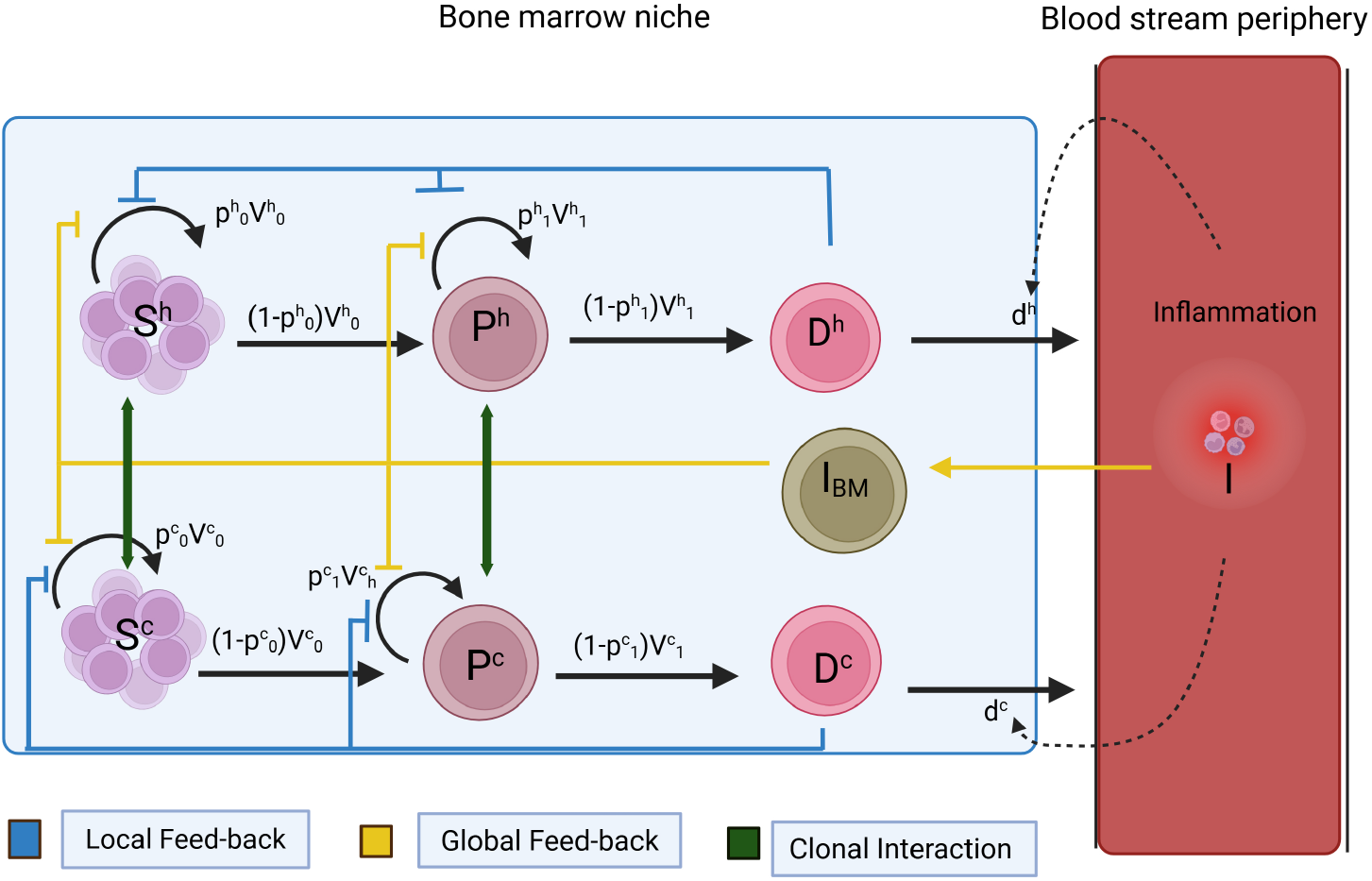
Schematic diagram of the two clonal global feedback inflammation model with clonal interaction term.

A simulation example illustrates the dynamic competition between a healthy hematopoietic population and a malignant clone, defined as a cell population that has acquired mutations disrupting selfrenewal feedbacks (high 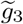). The simulation in Figure (10) shows that the healthy population declines as the malignant clone grows, causing MDS within 100 days. The system starts with healthy stem, progenitor, and differentiated cells. However, leukemic stem cells start to suppress healthy progenitor and differentiated cells, resulting in the disruption of normal hematopoiesis and malignant clone dominance.

**Figure 10.**
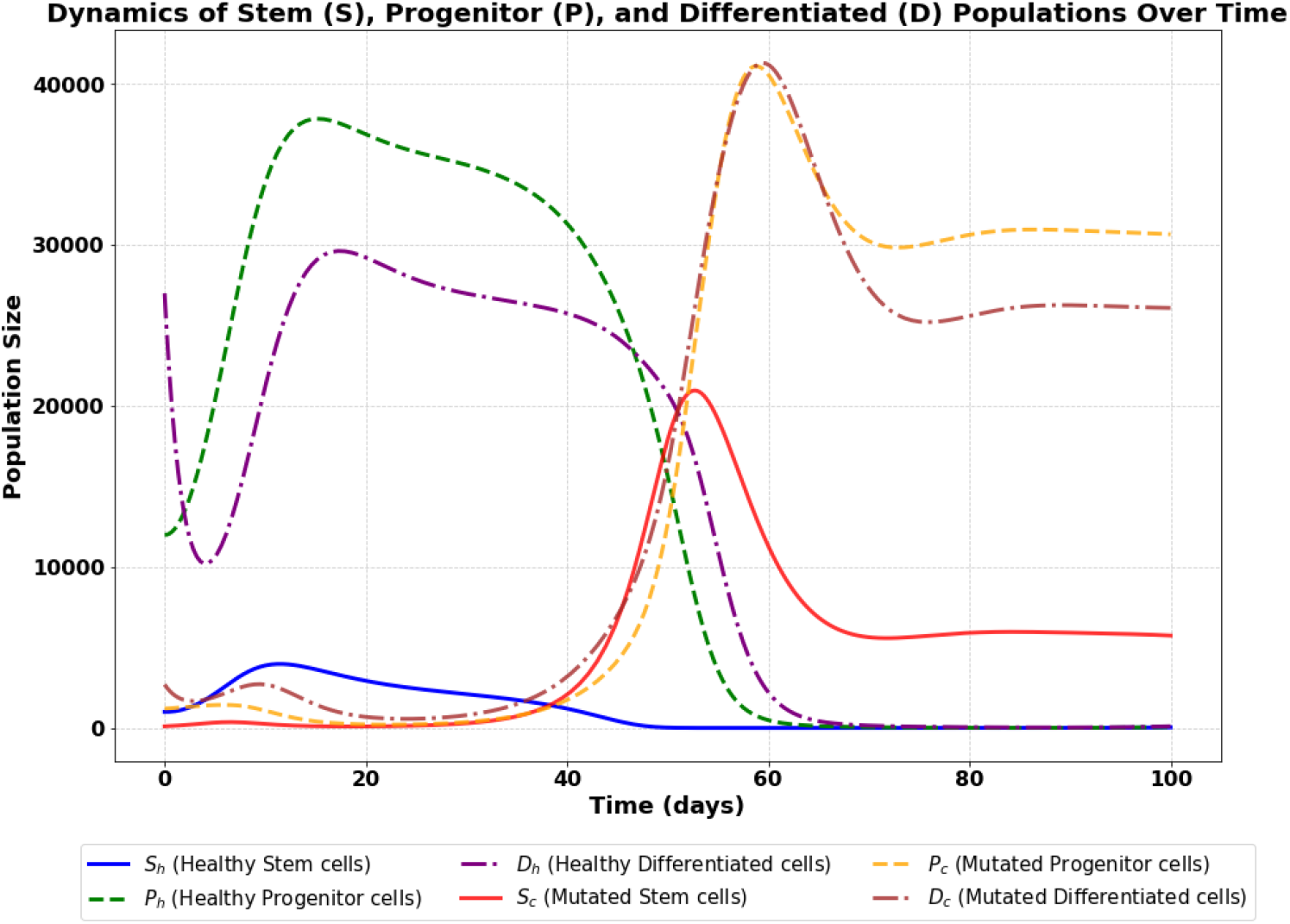
Progression of MDS dynamics between healthy and mutated cell populations under inflammation.

The corresponding sensitivity analysis (Figs. 4 and 5 in SI) quantifies how variations in model parameters influence the behavior of healthy and cancerous populations. For healthy cells, the most influential parameters are 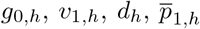, and *K*_*h*_, which regulate self-renewal, differentiation, and cell death.

These ensure hematopoietic homeostasis by maintaining a balance between cell production and clearance. In particular, *g*_0*h*_ governs how terminal cell density feeds back on stem cell self renewal, a mechanism essential during stress or infection. The differentiation rate *v*_1,*h*_ and the death rate *d*_*h*_ determine how efficiently progenitors become functional immune cells and are cleared.

In contrast, malignant populations are most sensitive to 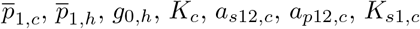, and *K*_*p*1,*c*_. Notably, the proliferation rate of malignant progenitors 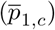 dominates, reflecting their aggressive clonal expansion. Parameters like 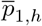 and *g*_0,*h*_, although linked to healthy cells, affect malignant expansion through niche competition and shared feedback mechanisms. Larger leukemic carrying capacities enable malignant clones to grow despite competition, especially when healthy populations are suppressed.

The sensitivity analysis results demonstrate how the interplay between mutated feedback loops, competition dynamics, and key regulatory parameters leads to the dominance of malignant clones and the progression from clonal hematopoiesis to MDS. The analysis highlights that progenitor proliferation and feedback sensitivity are the most critical drivers of this transition, providing clear targets for future therapeutic modeling and intervention.

## 4 Discussion

Bone marrow plays a critical role in the regulation of inflammation, being responsible for the generation of immune cells [1]. Several lines of evidence suggest a bidirectional interplay between clonal hematopoiesis and inflammation, since clonal cell populations fuel inflammation whereas inflammation enables mutated clones to outcompete wild-type clones and expand [60]. Inflammatory mediators, such as cytokines or pathogen-derived molecules, drive proliferation and differentiation of HSC towards myeloid lineage, which results in the functional decline of HSC and their attrition [1]. Additionally, unresolving chronic inflammation and sustained myeloid differentiation of progenitor cells drive pathologies, such as cardiovascular disease [61]. In a similar manner, several studies suggest that mutated myeloid cells in subjects with CHIP have pro-inflammatory properties, resulting in increased frequency and severity of chronic inflammatory disorders such as myocardial infarction or stroke [62, 63]. On the other hand, clonal cells are resistant to inflammation and are able to maintain their functional properties [21, 64]. Specifically, disruption of small intestinal barrier and bacterial translocation takes place in Tet2 deficient mice, which further drives pre-leukemic myeloproliferation through IL-6 production [64]. In the case of Dnmt3a deficiency, mutated clones were able to expand during mycobacterial infection, suggesting that these cells are resistant to IFN-*γ* mediated suppression of HSC function [21].

We took advantage of these clinical and experimental observations to develop a model that could integrate inflammation in the classical models of hematopoiesis that have been developed so far. We provide a thorough mathematical framework to examine the complex dynamics of hematopoiesis under local or global feedbacks with chronic inflammation and clonal hematopoiesis. We have shown the development from healthy hematological states to clonal disorders like MDS and AML by integrating inflammatory signal-influenced feedback mechanisms. Emergency myelopoiesis and its transition to malignancy were simulated using inflammation-dependent parameters, demonstrating the distinct vulnerability of HSCs and progenitor cells to inflammatory stimuli.

Furthermore, our model underscores the importance of parameter tuning and threshold effects in the transition between hematopoietic states. For example, higher values of inflammation-dependent parameters (*K, g*_2_ and *g*_3_) were found to destabilize the system, pushing it toward pathological states. Conversely, stabilizing factors like higher values of *g*_2_ disease progression suggest potential avenues for therapeutic intervention.

Sensitivity analysis key findings include the identification of critical parameters that drive the system’s behavior under varying conditions. Sensitivity analysis revealed that the proliferation rates of progenitor cells (*p*_1,*h*_, *p*_1,*c*_) and carrying capacity terms are pivotal in modulating the dynamics of both healthy and cancerous cell populations. These insights emphasize the role of feedback mechanisms and clonal competition in determining the fate of the hematopoietic system. Notably, the model highlighted how pathological inflammation can amplify the dominance of mutated clones over healthy populations, creating a self-reinforcing loop that promotes disease progression.

Nevertheless, the model operates under certain assumptions that warrant further discussion. The use of Hill functions to model feedback effects simplifies complex biological signaling cascades and may not fully capture time-delayed or nonlinear interactions present in vivo. Additionally, we assume homogeneous stem and progenitor populations, whereas the biological reality is far more heterogeneous, including quiescent, activated, and lineage-biased subpopulations. Future refinements could involve incorporating stochastic elements or age-structured compartments to better reflect this diversity. Furthermore, inflammation is treated under a quasi-steady-state approximation, which may overlook cytokine pulse dynamics and short-term perturbations that are relevant in chronic infection or treatment response. Comparing our approach to previous modeling efforts, we extend the classical models [28, 32, 35], which focused predominantly on lineage differentiation and mutation accumulation, by integrating inflammation as both a dynamic regulator and a disease-driving mechanism. Unlike prior work that treated inflammatory effects as static or external perturbations, our model embeds inflammation within the feedback structure of hematopoiesis. This provides a more physiologically realistic depiction of how inflammation both arises from and alters hematopoietic output, forming a feedback loop that favors mutated clones.

Looking forward, future model iterations could incorporate spatial heterogeneity in the bone marrow niche or multi-lineage interactions including erythroid and lymphoid cells, which are also susceptible to inflammatory regulation. Incorporating stochastic mutations and niche competition via agent-based modeling could further enrich the realism of clonal dynamics. Ultimately, integrating this modeling framework with longitudinal patient data, inflammation biomarkers, and single-cell transcriptomic profiles could enable precision hematology by identifying early inflection points in disease progression and tailoring anti-inflammatory or differentiation-based therapies accordingly.

## 5 Conclusion

This study provides a new mathematical paradigm for understanding inflammation and clonal hematopoiesis. Our observations show that inflammatory feedback loops drive clonal selection and proliferation, giving mechanistic insights into MDS and AML progression. The sensitivity analysis identified critical factors that could be biomarkers or therapeutic targets, demonstrating the power of mathematical modeling to comprehend complicated biological systems. Genetic mutations and microenvironmental heterogeneity should be added to the model in the future. Experimental validation of parameters and feedback mechanisms is necessary to improve model prediction. Combining this paradigm with clinical data could enable tailored disease progression and anti-inflammatory therapy predictions. This study increases our understanding of hematopoietic dynamics and lays the groundwork for developing new treatments to reduce chronic inflammation and clonal proliferation in hematological malignancies.

## Supporting information

SI_updated

## Acknowledgement

HH would like to thank Volkswagenstiftung for its support of the “Life?” program (96732). HH, CP, SV and IM acknowledge the support of the RIG-2023-051 grant from Khalifa University. Finally, HH would like to thank for the support of the UAE-NIH Collaborative Research grant AJF-NIH-25-KU.

HH also acknowledges Anna Maria Pappa and the University of Cambridge for providing access to BioRender, which was used to create the following figures: Graphical abstract; https://BioRender.com/ovrj6e3, Figure 1; https://BioRender.com/7azi1ek Figure 4; https://BioRender.com/83ntruv and Figure 9; https://BioRender.com/bvtpuxn

## Authors Contributions

Conceptualization; H.H; methodology: H,H, S.S, Y.J.U and I.M; software; S.S, Y.J.U; formal analysis; H.H, S.S, Y.J.U and I.M; writing—original draft preparation; H.H, S.S, Y.J.U and I.M; writing, reviewing and editing; H.H, S.S, Y.J.U, C.P and I.M; visualization; H.H, S.S, Y.J.U, C.P and I.M; supervision; H.H and I.M;

## Ethics approval and consent to participate

This study was conducted entirely through in silico analysis using computational modeling. No human or animal data were collected or utilized, and therefore ethical approval and informed consent were not required.

## Declaration of generative AI and AI-assisted technologies in the writing process

During the preparation of this work, the authors used ChatGPT/OpenAI in order to improve the language and correct the grammar of the text. After using this tool/service, the authors reviewed and edited the content as needed and take full responsibility for the content of the published article.

## Data and materials availability

All of the data supporting the results in this study are available within the paper and its supplementary information.

