## Supplementary material for "In Silico Investigation of the Role of Local and Global Inflammation-Driven Feedback in Myelopoiesis and Clonal Cell Expansion": SI_updated

May 2025

2.3 Model with two distinct populations (healthy and clonal) with dynamic Inflammation . . 10

**3 Global sensitivity analysis** **12**

**4 Initial conditions and parameter values** **13**

**5 Statistical analysis of the local feedback model** **14**

**6 Clonal global sensitivity analysis** **15**

### 1 Introduction

This study explores three mathematical models that describe the dynamics of stem cells, progenitor cells, and differentiated cells within the bone marrow environment. Beginning with a simplified hematopoiesis model, based on the framework proposed by Manciniak et al. [1] with a minor modification (considering decay only in the differentiated cell compartment), we first analyzed the system without inflammation and then introduced static inflammation.

Subsequently, the model was extended to capture the effects of chronic inflammation, focusing on how inflammation in the peripheral blood propagates to the bone marrow, leading to mutations that genetically alter the self-renewal rates of both stem and progenitor cells. Recognizing the complexity inherent in mathematical modeling, the study followed a structured approach: starting with a simple model to analyze key parameters and parameter ranges before progressing to more intricate systems.

The procedure adopted is as follows:

Model 1 comes from the classical hematopoiesis model proposed by Anna Marciniak et al [1]. Death of cells is assumed to occur only after maturation. In our case we assumed cell death can be influenced by static inflammation implying constant inflammation over a short period of time. The traditional assumption of regulatory feedback acting on the self-renewal rates of stem and progenitor cells due to the density of matured cells is maintained. Feedback functions are modeled using Hill functions, which are well suited for characterizing ligand-receptor interactions and their regulatory effects.

Model 2 is an improvement of model one with an added complexity. We incorporated the influence of chronic inflammation from the peripheral blood that leads to inflammation in the bone marrow, representing prolonged inflammatory states. This model seeks to investigate the behavior of hematopoiesis under sustained inflammatory conditions.

In Model 3, we separated the population of healthy normal cell populations and their clonal counterparts. Our intuition is to investigate the parameter regimes that lead to clonal cell population dominating the healthy hematopoietic cell population under external factors such as chronic inflammation.

Through comparative analysis of these models, the study aims to provide insight into the role of inflammation in clonal hematopoiesis, with implications for understanding disease progression and treatment strategies.

### 2 Model types

#### 2.1 Hematopoiesis model with local feedback (HMLF)

This section represents the first model. Here, we employed the classical model of hematopoesis and assumed that static inflammation is affected by a sudden change in the apoptotic rate. This behavior can be represented by a pertubed parameter $d$. By assuming that $S$, $P$ and $D$ represent the stem, progenitor and differentiated cell populations respectively, we can represent the dynamics of the populations as shown in figure 1 by the following odes'

$$\dot{S} = (2p_0 - 1)v_0 S \tag{2.1}$$

$$\dot{P} = 2(1 - p_0)v_0 S + (2p_1 - 1)v_1 P \tag{2.2}$$

$$\dot{D} = 2(1 - p_1)v_1 P - dD \tag{2.3}$$

where equation (2.1) describes the dynamics of the stem cell (S) population over time. Equation (2.2) represents the dynamics of the progenitor cell (P) population. Equation (2.3) captures the dynamics of the differentiated cell (D). $p_0, v_0$ represent the self renewal rate and division rate of stem cells respectively. The self renewal rate and division rate of progenitor cells are represented by $p_1, v_1$. During peripheral inflammation, signals such as tumor necrosis factor-alpha (TNF-$\alpha$) or interleukin-1 (IL-1) can diffuse into the bone marrow, increasing apoptosis rates in a manner proportional to the inflammatory burden.

A key aspect of hematopoiesis is its tight regulation through feedback mechanisms. In this model, negative feedback is mediated by the density of differentiated cells, which influences the self-renewal probabilities of stem and progenitor cells. This regulation is captured using Hill functions, which are widely used to describe sigmoidal biological responses. The feedback functions are given as:

$$p_0 = \frac{\bar{p_0}}{1 + g_0 D} \tag{2.4}$$

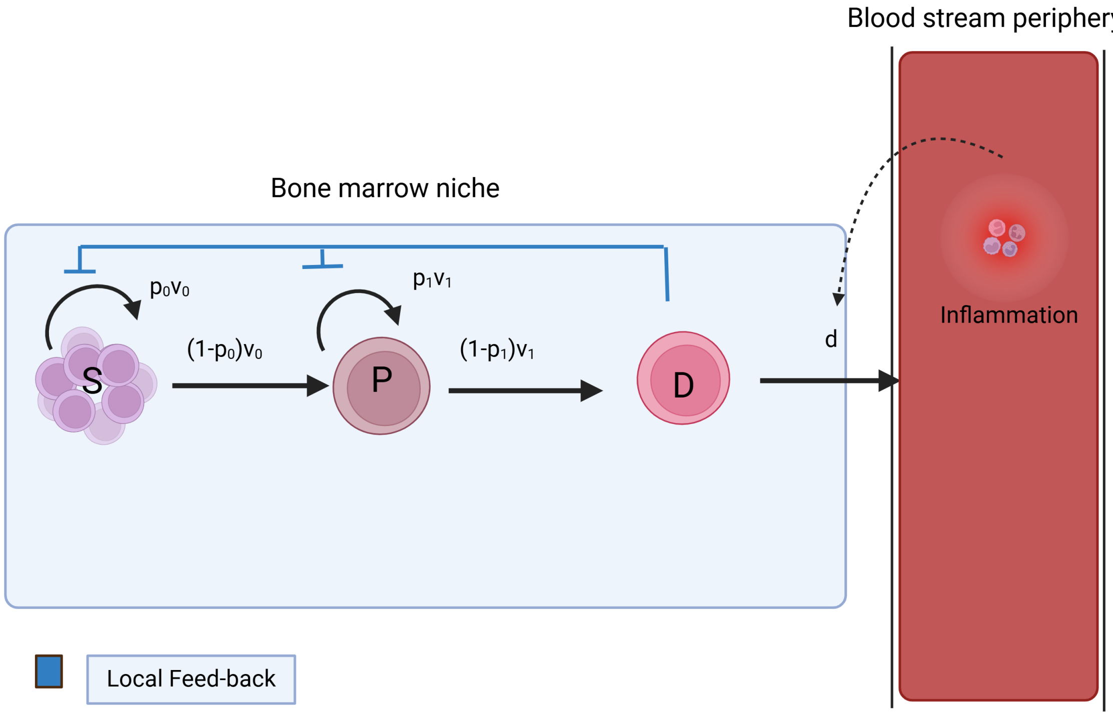

Figure 1: Schematic diagram of hematopoiesis model with static inflammation

$$p_1 = \frac{\bar{p_1}}{1 + g_1 D} \tag{2.5}$$

where $\bar{p_0}$ and $\bar{p_1}$, are the maximum possible feedback values for stem cell and progenitor cell self-renewal rates, respectively. $g_0$ and $g_1$ represent the sensitivity of the negative feedback mechanisms on stem and progenitor cell self-renewal. $\bar{p_0}$, $\bar{p_1} \in (0,1)$, with values closer to 0 indicating minimal inhibition and values closer to 1 indicating strong inhibition. The feedback mechanism reflects the biological regulation within the bone marrow, where the secretion of cytokines(small signaling proteins) modulates hematopoiesis. Cytokines are produced in response to the density of differentiated cells, influencing both stem and progenitor cell behavior.

Hill functions effectively model this regulatory process, as they account for the saturation effect implying that at high concentrations of differentiated cells, the feedback mechanism reaches a maximum inhibitory effect.

##### 2.1.1 Parameter estimation of local feedback model

At steady state, the equation for stem cells gives

$$\dot{S} = (2p_0 - 1)v_0 S = 0 \tag{2.6}$$

which implies $p_0 = \frac{1}{2}$

Similarly, we have from equation (2.2) that

$$\bar{P} = \frac{2(1-p_0)v_0\bar{S}}{(1-2p_1)v_1} \implies \frac{\bar{P}}{\bar{S}} = \frac{2(1-p_0)v_0}{(1-2p_1)v_1} \tag{2.7}$$

Furthermore, substituting $p_0 = \frac{1}{2}$ into equation (2.7)gives

$$\bar{P} = \frac{v_0\bar{S}}{(1-2p_1)v_1} \tag{2.8}$$

Finally, for the steady state of the differentiated cells, we get from (2.3) that

$$\bar{D} = \frac{2(1-p_1)v_1\bar{P}}{d} \tag{2.9}$$

which also implies

$$\bar{P} = \frac{d\bar{D}}{2(1-p_1)v_1} \tag{2.10}$$

Considering negative feedback on the model, we can set, for simplicity, $g_0 = g_1 = g$. Then we have

$$(2p_0 - 1)v_0\bar{S} = 0 \implies p_0 = \frac{1}{2} \implies \frac{\bar{p_0}}{1+g\bar{D}} = \frac{1}{2} \tag{2.11}$$

which simplify to

$$\bar{D} = \frac{2\bar{p_0} - 1}{g} \tag{2.12}$$

Hence by substituting (2.12) into (2.10), we get

$$\bar{P} = \frac{d\bar{D}}{2(1-p_1)v_1} = \frac{d}{gv_1}\frac{(2\bar{p_0} - 1)}{2(1-p_1)} \tag{2.13}$$

Next we calculated the ratio of progenitor over stem cells which serves as a criterion for the disease stage analysis:

$$\frac{\bar{P}}{\bar{S}} = \frac{2(1-p_0)v_0}{(1-2p_1)v_1} = ... = \frac{2v_0}{v_1}\frac{1+gD-\bar{p_0}}{1+gD-2\bar{p_1}} \tag{2.14}$$

**Normal condition**: physiological bone marrow tissues typically have plenty of diffrentiated cells, i.e. $D >>> 1$:

$$\frac{\bar{P}}{\bar{S}} \approx \frac{2v_0}{v_1} \geq 10 \implies v_0 \geq 5v_1. \tag{2.15}$$

The latter is a very important relationship since it allows us to calibrate the proliferation rates between stem cell and progenitors. Also it recapitulates our knowledge that the stem cell proliferative potential is much larger than the progenitor one.

**Pathological condition**: During infections, there is a high demand for differentiated immune cells, such as monocytes, and neutrophils. Therefore, a short-term depletion of differentiated cells can occur

$$D \longrightarrow 0$$

and using the Taylors expansion on equation (2.14), the ratio becomes

$$\frac{\bar{P}}{\bar{S}} \approx \frac{2v_0}{v_1}\left(\frac{\bar{p_0}-1}{2\bar{p_1}-1} + \frac{\bar{p_0}-2\bar{p_1}}{(1-2\bar{p_1})^2}D\right) \tag{2.16}$$

In this case the ratio is modulated according to the baseline self-renewal rates of stem and progenitor cells.

The parameters $\bar{p_0}$ and $\bar{p_1}$ can change from their physiological values only by means of mutations. Therefore, we would like to investigate what is the impact of such mutations in the robustness of myelopoiesis. If

$$\bar{p_0} > 2\bar{p_1} \implies \frac{\bar{P}}{\bar{S}} \downarrow \text{as D} \downarrow$$

which implies during inflammation (reducing differentiated cells) there is a possibility of transition from normal to MDS or even to AML.

On the other hand if

$$\bar{p_0} \leq 2\bar{p_1} \implies \frac{\bar{P}}{\bar{S}} \uparrow \text{as D} \downarrow$$

which implies that the bone marrow remains in its Normal state.

Similarly,

From equation (2.1) and (2.4) we have at steady state,

$$\dot{S} = 0 \implies g_0 = \frac{1}{\bar{D}} \tag{2.17}$$

Let

$$\frac{\bar{D}}{\bar{P}} = \lambda_{DP} \tag{2.18}$$

represent the ratio of differentiated cells to that of progenitors at steady state,

We now have

$$\frac{\bar{D}}{\bar{P}} = \left(\frac{2v_1}{d}\right)\left(1 - \frac{\bar{p_1}}{1 + \dfrac{g_1}{g_0}}\right) \tag{2.19}$$

Next by letting $g' = \frac{g_1}{g_0}$ we have

$$\frac{2v_1}{d}\frac{1 + g_1' - \bar{p_1}}{(1 + g_1')} = \lambda_{DP} \tag{2.20}$$

resulting in

$$\bar{p_1} = 1 + g_1' - \frac{d\lambda_{DP}}{2v_1}(1 + g_1')$$

and finally we have

$$\bar{p_1} = (1 + \frac{g_1}{g_0})(1 - \frac{d\lambda_{DP}}{2v_1}) \tag{2.21}$$

from equation 2.21 we can see that since $g_1$ and $g_0$ are positive and $\bar{p_1} \geq 0$, then

$$1 - \frac{d\lambda_{DP}}{2v_1} \geq 0 \tag{2.22}$$

which implies that

$$1 \geq \frac{d\lambda_{DP}}{2v_1} \implies \frac{v_1}{d} \geq \frac{\lambda_{DP}}{2}. \tag{2.23}$$

Similarly, from equation 2.20, we see that $1 + \dfrac{g_1}{g_0} - \bar{p_1} \geq 0$ implying that $\bar{p_1} \leq 1 + \dfrac{g_1}{g_0}$ which is trivial since $\bar{p_1} \in (0, 1)$, and $\dfrac{g_1}{g_0} \geq 0$

At steady state, we have

$$\frac{\bar{P}}{\bar{S}} = \lambda_{PS} \implies \bar{S} = \frac{1}{g_0\lambda_{DP}\lambda_{PS}} = (\lambda_{DP}\lambda_{PS})^{-1}\frac{1}{g_0} \tag{2.24}$$

since $\frac{\bar{D}}{\bar{P}} = \lambda_{DP}$ and $\bar{D} = \frac{1}{g_0}$ .

Therefore

$$\lambda_{PS} = \frac{v_0}{v_1}\frac{1 + \dfrac{g_1}{g_0}}{1 + \dfrac{g_1}{g_0} - 2\bar{p_1}} \tag{2.25}$$

The expression

$$1 + \frac{g_1}{g_0} - 2\bar{p_1} \geq 0 \tag{2.26}$$

must hold for a steady state ratio of progenitors to stem cells.

This implies $\bar{p_1} \leq \frac{1}{2} + \frac{1}{2}\frac{g_1}{g_0}$ if $g_1 \geq g_0$ and $\bar{p_1} < 1$ if $g_1 \leq g_0$, with the former always being true because

$$1 + g_1' - 2\bar{p_1} = \frac{v_0}{v_1\lambda_{PS}}(1 + \frac{g_1}{g_0})$$

$$\implies 2\bar{p_1} = (1 + \frac{g_1}{g_0}) - \frac{v_0}{v_1\lambda_{PS}}(1 + \frac{g_1}{g_0})$$

$$\implies \bar{p_1} = \frac{1}{2}(1 + \frac{g_1}{g_0})(1 - \frac{v_0}{v_1\lambda_{PS}}) \tag{2.27}$$

From 2.27 , we can deduce that

$$1 - \frac{v_0}{v_1\lambda_{PS}} \geq 0 \implies \frac{v_0}{v_1} \leq \lambda_{PS} \tag{2.28}$$

equating 2.21 and 2.27 we have

$$\left((1+\frac{g_1}{g_0})(1-\frac{d\lambda_{DP}}{2v_1})\right)=\frac{1}{2}(1+\frac{g_1}{g_0})(1-\frac{v_0}{v_1\lambda_{PS}})$$

and

$$v_1 = d\lambda_{DP} - \frac{v_0}{\lambda_{PS}} \tag{2.29}$$

#### 2.2 Hematopoiesis model with global feedback (HMGF)

Here, we studied the hematopoiesis model influenced by chronic Inflammation from the peripheral blood as shown in Figure (2). We assumed that inflammation in the peripheral blood leads to inflammation in the bone marrow. $\dot{S}$ $\dot{P}$ and $\dot{D}$ represent the dynamics of stem, progenitor and differentiated cell populations, respectively. $\dot{I}$ represent the dynamics of inflammation in the peripheral blood.$\dot{I_{BM}}$ represent the dynamics of inflammation in the bone marrow.

Parameters $\beta$, $a$, $d_1$, and $d_2$ play important roles in describing the dynamics of inflammation within both the bloodstream and bone marrow environments.

Parameter $a$ represents the rate at which differentiated cells contribute to impacting inflammation in the bloodstream. This parameter captures the extent to which cellular differentiation affects the chronic inflammatory levels. $d_1$ is the combined rate of inflammatory decay within the bloodstream, encompassing both natural clearance and the transport of inflammatory markers to the bone marrow.

Thus $d_1$ represent systemic clearance of inflammatory markers from the bloodstream, which may be transported into or otherwise influence inflammation into the bone marrow. $\beta$ represents the fraction rate of $d_1$ that corresponds to the portion of inflammatory markers from the bloodstream that influence inflammation in the bone marrow.

Finally, $d_2$ represents the rate of decay of inflammation in the bone marrow. This decay results in cues that cause negative feedback on stem and progenitor cell self renewal rates.Thus, it captures both the natural resolution of inflammation within the bone marrow and the strength of negative feedback on hematopoietic stem and progenitor cells. These negative feedbacks resulting from the localized bone marrow inflammation is assumed to inhibit self renewal rates of the cells.

$$\dot{S} = (2p_0 - 1)v_0 S \tag{2.30}$$

$$\dot{P} = 2(1 - p_0)v_0 S + (2p_1 - 1)v_1 P \tag{2.31}$$

$$\dot{D} = 2(1 - p_1)v_1 P - d(I)D \tag{2.32}$$

$$\dot{I} = ad(I)D - d_1 I \tag{2.33}$$

$$\dot{I_{BM}} = \beta I - d_2 I_{BM} \tag{2.34}$$

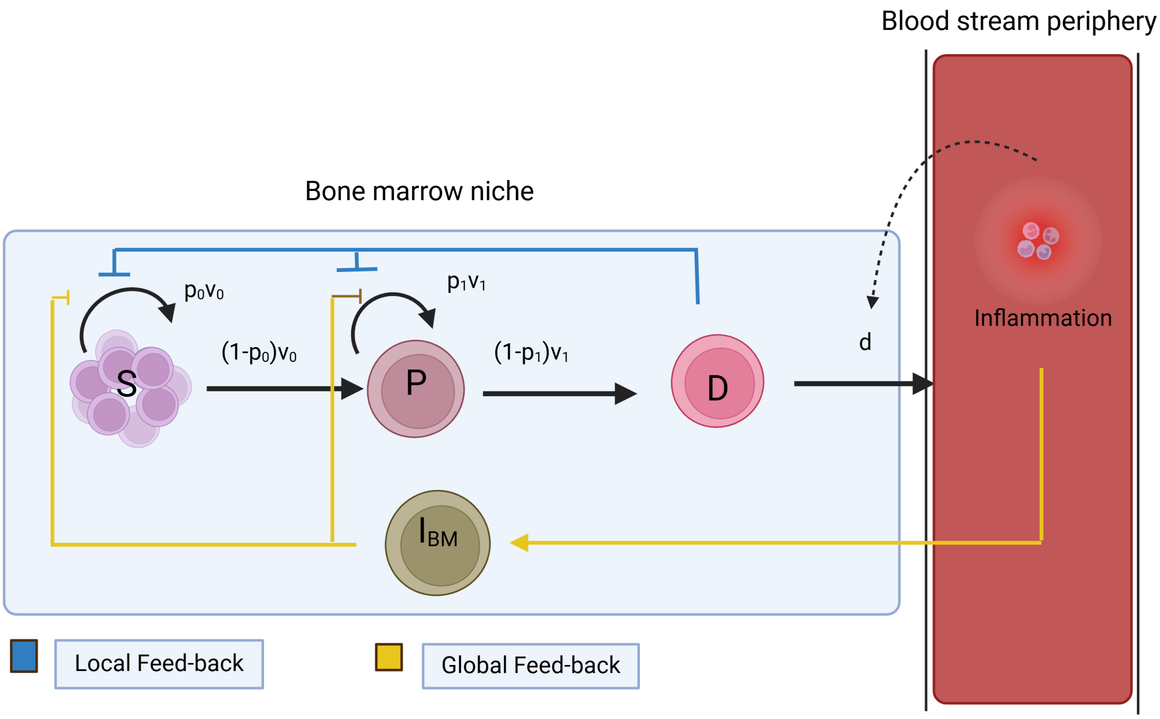

Figure 2: Schematic diagram of the global feedback inflammation model.

The negative feedback regimes are now influenced by inflammation in the bone marrow. We represent the negative feedbacks mathematically as follows.

$$p_0 = \frac{\bar{p_o}}{1 + g_0 D + g_2 I_{BM}} \tag{2.35}$$

and

$$p_1 = \frac{\bar{p_1}}{1 + g_1 D + g_3 I_{BM}} \tag{2.36}$$

where $g_0$ and $g_1$ represent degree of sensitivity of negative feedbacks on stem cells and preogenitor cell self renewal rate respectively due to density of matured cells. Parameters $g_2$ and $g_3$ represent degree of influence of inflammation in bone marrow that contributes to negative feedback or inhibition of self renewal of stem cells and that of progenitors.

##### 2.2.1 HMGF model reduction under quasi-steady state assumption

Under the assumption of a quasi-steady state for inflammation, we set at steady state,

$$I_{BM} = \frac{\beta}{d_2} I \tag{2.37}$$

and

$$I = \frac{ad(I)D}{d_1}. \tag{2.38}$$

Now assuming that the outflux of the terminal cells is governed by a monotonically increasing function

$$d(I) = dK \frac{I+1}{I+K} \tag{2.39}$$

where $K$ controls the sensitivity of cell death due to inflammation. When $K$ is small(supposedly at minimum value of 1), $d(I)$ remains close to its basal value, ensuring homeostasis

From equation (2.33), we can represent $I$ as

$$I = \frac{ad(I)D}{d_1} = \frac{adK}{d_1} \frac{1+I}{I+K} D \tag{2.40}$$

Assuming

$$\frac{ad}{d_1} = \frac{1}{a'} \tag{2.41}$$

equation (2.40) becomes

$$a'I(I+K) = K(1+I)D \implies a'I^2 + a'KI = DK + DIK \iff a'I^2 + (a'K - DK)I - KD = 0 \tag{2.42}$$

Solving the quadratic equation, we have

$$I = \frac{DK - a'K \pm \sqrt{(a'K - DK)^2 + 4a'DK}}{2a'} \tag{2.43}$$

Taking $D \gg 1$, the term inside the square root, $(a'K - DK)^2 + 4a'DK$, can be expanded :

$$(a'K - DK)^2 = (DK)^2 - 2a'K^2 D + (a'K)^2. \tag{2.44}$$

Thus the radicand of equation (2.43) becomes

$$(a'K - DK)^2 + 4a'DK = (DK)^2 - 2a'K^2 D + (a'K)^2 + 4a'KD. \tag{2.45}$$

Factoring out $D^2$, we get:

$$(a'K - DK)^2 + 4a'DK = (DK)^2 \left[1 - \frac{2a'}{D} + \frac{(a')^2}{D^2} + \frac{4a'}{DK}\right]. \tag{2.46}$$

For $D \gg 1$, terms proportional to $1/D^2$ or smaller become negligible compared to terms proportional to $1/D$. Using a first-order Taylor expansion for $\sqrt{1+x}$:

$$\sqrt{1+x} \approx 1 + \frac{x}{2} \quad (\text{when } |x| \ll 1), \tag{2.47}$$

where:

$$x = -\frac{2a'}{D} + \frac{4a'}{DK}. \tag{2.48}$$

The square root becomes:

$$\sqrt{(a'K - DK)^2 + 4a'DK} \approx DK \left[1 - \frac{a'}{D} + \frac{2a'}{DK}\right], \tag{2.49}$$

which simplifies to:

$$\sqrt{(a'K - D)^2 + 4a'D} \approx DK - a'K + 2a'.$$

Substituting the approximation for the square root back into the quadratic formula:

$$I = \frac{DK - a'K + \sqrt{(a'K - DK)^2 + 4a'DK}}{2a'},$$

we get:

$$I \approx \frac{DK - a'K + (DK - a'K + 2a')}{2a'}. \tag{2.50}$$

Simplifying the numerator:

$$I \approx \frac{2DK - 2a'K + 2a'}{2a'} \tag{2.51}$$

and factoring out 2:

$$I \approx \frac{DK - a'K + a'}{a'} \tag{2.52}$$

Finally:

$$I \approx \frac{DK}{a'} - K + 1. \tag{2.53}$$

From (2.53) , we see that for large $D$, the dominant term in $I$ is $\frac{DK}{a'}$ which scales linearly with $K$. From (2.39), by assuming that the range of

$$\frac{I+1}{I+K} \in (\frac{1}{K}, 1)$$

we have

$$d(I) \in (d, dK)$$

As such, equation (2.34) becomes

$$\dot{D} = 2(1-p_1)v_1P - d(I)D = 2(1-p_1)v_1P - dKD \tag{2.54}$$

and the complexity of the model is reduced to three ODEs as follows

$$\dot{S} = (2p_0 - 1)v_0S \tag{2.55}$$

$$\dot{P} = 2(1-p_0)v_0S + (2p_1 - 1)v_1P \tag{2.56}$$

$$\dot{D} = 2(1-p_1)v_1P - dKD \tag{2.57}$$

Next, due to lack of sufficient data on the exact population/concentration of inflammation in the bone marrow, we now transformed the negative feed backs to eliminate the bone marrow inflammation ($I_{BM}$) term. We assume that $d(I)$ varies within the range $(d, dK_{max})$ where $K \in (1, K_{max})$. Here, 1 represents the minimum value for the parameter $K$ with an unknown maximum value represented as $K_{max}$. Hence, at steady state,

$$I = \frac{ad(I)D}{d_1} \implies \frac{adD}{d_1} \leq I \leq \frac{adK_{max}D}{d_1} \tag{2.58}$$

substituting into $I_{BM}$ (equation 2.37), we have

$$\frac{a\beta dD}{d_2 d_1} \leq I_{BM} \leq \frac{a\beta dK_{max}D}{d_2 d_1} \tag{2.59}$$

Therefore, $I_{BM}$ equals for different values of $K$:

$$I_{BM} = g_{new}DK \tag{2.60}$$

with

$$g_{new} = \frac{a\beta d}{d_2 d_1}$$

Thus, equations 2.35 and 2.36 becomes:

$$p_0 = \frac{\bar{p_o}}{1 + g_0 D + \tilde{g_2} KD} \tag{2.61}$$

and

$$p_1 = \frac{\bar{p_1}}{1 + g_1 D + \tilde{g_3} KD} \tag{2.62}$$

Where

$$\tilde{g_2} = g_{new} g_2 = \frac{g_2 a\beta d}{d_2 d_1} \tag{2.63}$$

and

$$\tilde{g_3} = g_{new} g_3 = \frac{g_3 a\beta d}{d_2 d_1} \tag{2.64}$$

#### 2.3 Model with two distinct populations (healthy and clonal) with dynamic Inflammation

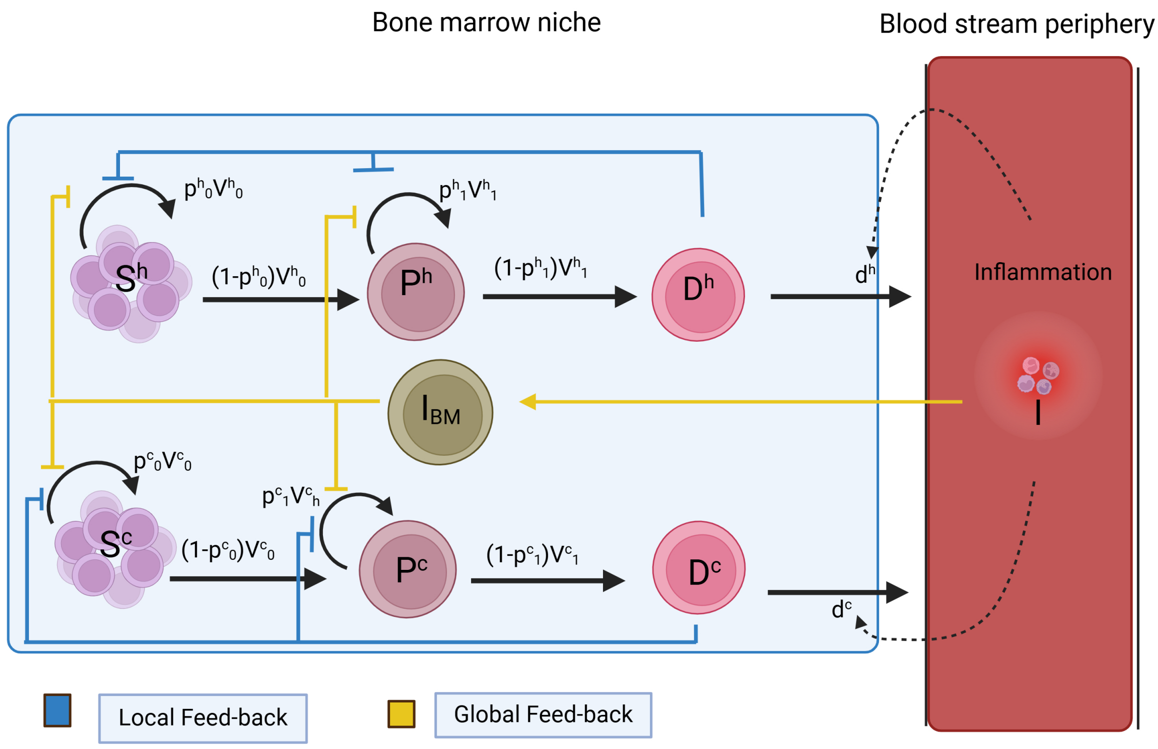

Figure 3: Schematic diagram of the two clonal global feedback inflammation model.

To gain a comprehensive understanding of the interactions and competitive behaviors between two distinct cell populations, a healthy cell population and a cancerous cell population, we employed a competition-based modeling approach. This framework was specifically designed to elucidate how the growth and proliferation of one population influences the dynamics of the other. By integrating competition terms into the kinetic equations, the model captures the inhibitory effects exerted by each population on the other, thereby simulating the complex biological interactions that arise in such systems. This approach allows for the exploration of critical parameters, such as growth rates, carrying capacities, and interpopulation competition coefficients, providing valuable insights into the dynamic balance between healthy and cancerous cell populations. Furthermore, the model serves as a basis for understanding the resource limitations and competitive pressures that govern their coexistence or dominance under various conditions.

The model is similar to the one presented in Section 2.2, which addresses global feedback with chronic inflammation, and its equations are as follows:

$$\dot{S}_h = (2p_{0,h} - 1 - a_{S,h}\frac{S_c}{K_{Sh}})v_{0,h}S_h \tag{2.65}$$

$$\dot{P}_h = 2(1 - p_{0,h} + a_{S,h}\frac{S_c}{K_{Sh}})v_{0,h}S_h + (2p_{1,h} - 1 - a_{P,h}\frac{P_c}{K_{Ph}})v_{1,h}P_h \tag{2.66}$$

$$\dot{D}_h = 2(1 - p_{1,h} + a_{p,h}\frac{P_h}{K_{Pc}})v_{1,c}P_h - d_h(I_h)D_h \tag{2.67}$$

$$\dot{S}_c = (2p_{0,c} - 1 - a_{S,c}\frac{S_h}{K_{Sc}})v_{0,c}S_c \tag{2.68}$$

$$\dot{P}_c = 2(1 - p_{0,c} + a_{S,c}\frac{S_h}{K_{Sc}})v_{0,c}S_c + (2p_{1,c} - 1 - a_{P,c}\frac{P_h}{K_{Pc}})v_{1,c}P_c \tag{2.69}$$

$$\dot{D}_c = 2(1 - p_{1,c} + a_{p,c}\frac{P_c}{K_{Pc}})v_{1,c}P_c - d_c(I_c)D_c \tag{2.70}$$

The subscripts $h$ and $c$ represents the healthy and cancerous populations, respectively.The parameters $a_S$ and $a_P$ denote the competition coefficients for the stem cell and progenitor populations, while $K_S$ and $K_P$ represent their respective carrying capacities. A simulated example in figure 4 of clonal evolution shows the dynamic interaction between a healthy cell population and a malignant clone. In this scenario, the malignant clone grows as the healthy cell population declines. MDS develops within 100 days due to this population dynamics shift. Cancerous clones gradually dominate the model, showing how clonal proliferation affects hematopoiesis and advances disease. More precisely, initially, the healthy populations dominate, with stem cells, progenitor cells, and differentiated cells maintaining stable levels. However, the emergence and expansion of cancerous stem cells initiate a clonal evolution, driving significant increases in cancerous progenitors and differentiated cells over time. As cancerous populations proliferate, healthy progenitors and differentiated cells experience a rapid decline, reflecting the disruption of normal hematopoiesis. Approximately halfway through the simulation, cancerous cells dominate, a hallmark of MDS, leading to reduced functional hematopoietic output. This visualization underscores the competitive dynamics between healthy and cancerous cell populations, highlighting the gradual onset and establishment of MDS.

### 3 Global sensitivity analysis

Random Sampling - High Dimensional Mathematical Representation (RS-HDMR) is crucial for comprehending the influence of intricate model outputs on input variables. This method assesses all input components concurrently across their whole range of values, unlike local sensitivity analysis, which examines minor variations in input parameters around a particular point. This comprehensive method positively influences models with non-linear input interactions. The principal benefit of RS-HDMR is its capacity to attribute model output uncertainty to the uncertainty of input factors. This aids in identifying the inputs that are most crucial and require careful monitoring or management. It also aids in prioritizing resources to comprehend and eradicate uncertainty in essential inputs [2]. RS-HDMR is essential in environmental modeling, engineering, and economics, where models display complex non-linear properties [2–5]. RS-HDMR improves the precision, dependability, and decision-making capabilities of models by measuring the impact of inputs on output variability. Typically, local sensitivity studies do not consider interactions among input variables; however, RS-HDMR can illustrate the influence of alterations in one variable on another. Comprehending and addressing the uncertainty inherent in model predictions is crucial for dependable and resilient modeling, as evidenced by RS-HDMR [2]

RS-HDMR, akin to the Sobol global sensitivity analysis method, disaggregates a model's output into components of increasing dimensionality [6].

$$f(x) = f_0 + \sum_{i=1}^{n} f_i(x_i) + \sum_{i<j}^{n} f_{ij}(x_i, x_j) + \ldots + f_{12\ldots n}(x) \tag{3.1}$$

where:

$$f_0 = \int f(x)dx \tag{3.2}$$

$$f_i(x_i) = \int f(x)dx_i - f_0 \tag{3.3}$$

$$f_{ij}(x_i, x_j) = \int f(x)dx_{ij} - f_i(x_i) - f_j(x_j) - f_0 \tag{3.4}$$

and $x = [x_1, x_2, \ldots, x_n]$ is the input of parameters whose values change; $dx_i$ is the product $dx_1 dx_2 \ldots dx_n$; $dx_{ij}$ is the same product without $dx_i$ and $dx_j$. The decomposition, known as analysis of variance (ANOVA), generates distinct and independent terms and uncovers both primary and secondary effects to represent output using methods based on variance. The total variance $D$ is [7]:

$$D = \int f^2(x)dx - f_0^2 \tag{3.5}$$

Further, the partial variances $D_{ij\ldots}$ are calculated by:

$$D_i = \int f_i^2(x_i)dx_i \tag{3.6}$$

$$D_{ij} = \int\int f_{ij}^2(x_i, x_j)dx_i dx_j \tag{3.7}$$

The sensitivity indices are:

$$S_{i1,\ldots,is} = \frac{D_{i1,\ldots,is}}{D}, \quad 1 \le i_1 < \ldots < i_s \le n \tag{3.8}$$

Thus, all terms add up to 1:

$$\sum_{i=1}^{n} S_i + \sum_{i<j} S_{ij} + \ldots + S_{12\ldots n} = 1 \tag{3.9}$$

The principal effect of the input variable $x_i$ on the output is ascertained by the first-order sensitivity index $S_i$, while the influence of both $x_i$ and $x_j$ on the output is represented by the second-order sensitivity indices, and so on. Every model outcome.

### 4 Initial conditions and parameter values

For the local feedback model equations, the initial population values of $S$, $P$ and $D$ are set to 1000, 12000 and 27000, respectively [8]. In the global feedback inlammation model, the initial conditions for $I$, and $I_{BM}$ are assumed to be equal to 280, and 2200, respectively. Table 1 shows the nominal values of both model parameters.

Table 1: Nominal values of the model parameters

| Parameter | Value | Reference |
|---|---|---|
| $v_0$ | 0.53 | [8] |
| $v_1$ | 0.49 | [8] |
| $g_0$ | $3.7{\cdot}10^{-5}$ | [8] |
| $g_1$ | $2.8{\cdot}10^{-5}$ | [8] |
| $d$ | 0.23 | [8] |
| $\bar{p_0}$ | 1 | [8] |
| $\bar{p_1}$ | 0.818 | [8] |
| $a$ | 0.9 | Assumed |
| $K$ | 5 | Assumed |
| $d_1$ | 20 | Assumed |
| $\beta$ | 0.8 | Assumed |
| $d_2$ | 0.1 | Assumed |
| $g_2$ | $0.5{\cdot}10^{-5}$ | Assumed |
| $g_3$ | $0.5{\cdot}10^{-5}$ | Assumed |

#### 5 Statistical analysis of the local feedback model

| | | | | Kruskal Wallis | Mann Whitney | | |
|---|---|---|---|---|---|---|---|
| | Healthy | MDS | AML | All Classes | Healthy-MDS | Healthy- AML | MDS – AML |
| | Mean ± SD | Mean ± SD | Mean ± SD | p vs Bonf. p-values | p vs Bonf. p–value | p vs Bonf. p–value | p vs Bonf. p–value |
| $g_1(\times 10^-5)$ | 2.95 ± 1.32 | 3.73 ± 2.05 | 4.67 ± 2.30 | 0.001**/0.004 | 0.054/0.983 | 0.000***/0.004 | 0.021*/0.381 |
| $v_0$ | 0.54 ± 0.25 | 0.51 ± 0.20 | 0.37 ± 0.19 | 0.000***/0.002 | 0.387/1.000 | 0.000***/0.007 | 0.001**/0.012 |
| $v_1$ | 0.33 ± 0.10 | 0.56 ± 0.22 | 0.65 ± 0.18 | 0.000***/0.000*** | 0.000***/0.000*** | 0.000***/0.000*** | 0.041*/0.742 |
| $d$ | 0.17 ± 0.04 | 0.32 ± 0.12 | 0.46 ± 0.14 | 0.000***/0.000*** | 0.000***/0.000*** | 0.000***/0.000*** | 0.000***/0.000*** |
| $p_1$ | 0.75 ± 0.14 | 0.68 ± 0.20 | 0.42 ± 0.26 | 0.000***/0.000*** | 0.034*/0.611 | 0.000***/0.000*** | 0.000***/0.000*** |
| $p < 0.05; **-p < 0.01$ ; $***-p < 0.001$ | | | | | | | |

Table 2: Statistical Analysis of Parameter Differences Across Healthy, MDS, and AML Groups (Kruskal-Wallis and Mann-Whitney Tests with Bonferroni Correction

### 6 Clonal global sensitivity analysis

This section, a more complex approach to the model is studied in which the cell population is divided into two distinct populations: healthy cells and clonal cells, as illustrated in figure 3. Clonal cells are defined as cells that originate from a common ancestor and therefore share identical genetic mutations or alterations. The primary objective of this work is to design the model in a way that allows investigation into the parameters that would cause clonal cells to dominate over a long period, ultimately leading to the extinction of healthy cells. Nominal values for the clonal evolution are in Table 3.

By conducting sensitivity analysis, Figure 5 illustrates the sensitivity indices $S_i$ values of various model parameters influencing the dynamics of healthy $S, P$ and $D$ cell populations. Sensitivity indices provide a quantitative measure of how variations in each parameter affect the output, thereby highlighting their relative importance in the model. For the healthy cell populations, the sensitivity analysis revealed that the most influential parameters were $g_{0,h}, v_{1,h}, d_h, \bar{p_{1,h}}$ and $K_h$.

These parameters collectively regulate the essential processes of self-renewal, differentiation, proliferation, and cell death, all of which are critical for maintaining hematopoietic homeostasis. $g_{0,h}$ parameter represents the degree of sensitivity of negative feedback on stem cells due to terminal cell density. Its importance lies in its role as the foundation for hematopoietic regeneration. Variations in $g_{0,h}$ directly affect the capacity of stem cells to sustain the progenitor and differentiated cell populations, ensuring the replenishment of immune and functional cells in the body. A robust baseline growth rate is crucial, particularly during periods of stress or increased demand, such as infections or inflammation. $v_{1,h}$ parameter governs the rate at which progenitor cells transition to differentiated cells. Although the sensitivity index of $v_{1,h}$ is moderate compared to other parameters, it is a key determinant of the system's ability to produce functional immune cells. Efficient differentiation ensures that the hematopoietic system meets physiological demands, particularly under normal conditions where stable production of differentiated cells is essential for health.

The death rate of differentiated cells significantly influences the dynamics of the healthy system, as these cells represent the terminally differentiated populations performing critical functions like oxygen transport, infection defense, and coagulation. A balance between production and death is necessary to maintain homeostasis. An elevated $d_h$ could result in depletion of functional cells, whereas decreased $d_h$ could lead to overcrowding or resource constraints in the bone marrow. $p_{1,h}$ parameter is the most critical for progenitor dynamics, as it controls the rate at which progenitor cells proliferate. Progenitor cells act as a transitional stage between stem cells and differentiated cells, and their proliferation directly determines the availability of cells for differentiation. High sensitivity to $p_{1,h}$ highlights its significant role in maintaining the dynamic equilibrium between cell production and consumption. The carrying capacity $K_h$ represents the maximum number of cells that the system can support within the available resources and environmental constraints. This parameter is crucial because it sets the upper limit for population size, preventing overgrowth and maintaining the balance between cell populations. Changes in $K_h$ can significantly alter the dynamics by either facilitating growth or imposing limitations on the system, particularly under conditions of resource scarcity or stress.

Similarly, Figure 6 presents the sensitivity indices of model parameters affecting the cancerous $S, P$ and $D$ cell populations. The results highlight the critical parameters driving the clonal expansion and progression of cancerous populations. For the cancerous cell populations, the sensitivity analysis revealed that the most critical parameters include the maximum negative feedback on the self-renewal rates of healthy and cancerous progenitor cells $(p_{1,h}, p_{1,c})$, the degree of sensitivity of the negative feedback on the self renewal rate of healthy stem cells due to terminal cell density $(g_{0,h})$ the inflammation feedback response to cancerous cells ($(K_c)$ ), the interpopulation competition coefficients (as12,c, ap12,c), and the carrying capacities associated with competition $K_{s,1,c}, K_{p,1,c}$. These parameters play pivotal roles in shaping the interplay between healthy and cancerous populations, directly influencing clonal expansion and the progression of hematopoietic malignancies.$p_{1,h}$ is primarily associated with healthy progenitor cells, this parameter indirectly impacts the cancerous populations by influencing the availability of resources and niche space. A high proliferation rate in $P_h$ can suppress the expansion of $P_c$ through resource competition, emphasizing the interdependence of these populations.$p_{1,h}$ is the most influential parameter for cancerous progenitor cells, as it governs their aggressive proliferation. The heightened sensitivity of $p_{1,c}$ underscores its role in driving clonal expansion and the dominance of cancerous cells over healthy populations. The degree of sensitivity of the negative feedback on self-renewal rate of healthy stem cells due to terminal density $(g_{0,h})$ indirectly affects the cancerous population by determining the competitive balance in the hematopoietic niche. A higher $g_{0,h}$ enables healthy cells to maintain their dominance, thereby limiting the expansion of cancerous populations. Larger $K_c$ values favor cancerous cell dominance, especially in conditions of prolonged inflammation or reduced competition from healthy populations $a_{s12,c}$ and $a_{p12,c}$ coefficients quantify the inhibitory effects that healthy populations exert on the cancerous stem and progenitor cells. A higher value of $a_{a12,c}$ or $a_{p12,c}$ indicates stronger suppression of cancerous growth due to competition for niche space, signaling, or resources. The cancerous population's dynamics are highly sensitive to these competition terms, as their expansion depends on how much healthy populations can outcompete or suppress them. Lastly,$K_{s,1,c}$ and $K_{p,1,c}$ parameters are critical for modeling how cancerous populations adapt and expand despite competitive pressures. Larger $K_{s,1,c}$ or $K_{p,1,c}$ values indicate a higher tolerance of cancerous cells to competition, reflecting their ability to exploit resources or withstand inhibitory effects from healthy cells. The cancerous population's dynamics are intrinsically linked to the behavior of the healthy population. Parameters like $g_{0,h}, a_{s12,c}, a_{p12,c}, K_{s,1,c}, K_{p,1,c}$ directly represent the competitive interactions between these populations. A high $g_{0,h}$ or effective competition terms($a_{s12,c}$and $a_{p12,c}$) reduce the likelihood of cancerous stem cells achieving dominance. The proliferation of healthy progenitor cells ($p_h$, driven by $p_{1,h}$) constrains $p_c$ by limiting available resources and niche access. Cancerous carrying capacities ($K_{s,1,c}, K_{p,1,c}$) depend on the extent to which healthy populations utilize shared resources. A decline in healthy populations due to inflammation or other factors enables cancerous populations to expand into available space.

Figures 12, and 13 collectively provide a comprehensive sensitivity analysis for both healthy and cancerous cell populations. The dominance of parameters like $P_1$ in both populations emphasize their universal role in cell dynamics. However, subtle differences in sensitivity patterns highlight the distinct mechanisms driving healthy versus cancerous population dynamics. For example, the higher sensitivity of $P_1$ in cancerous progenitor cells underscores the aggressive proliferation characteristic of cancerous clones.

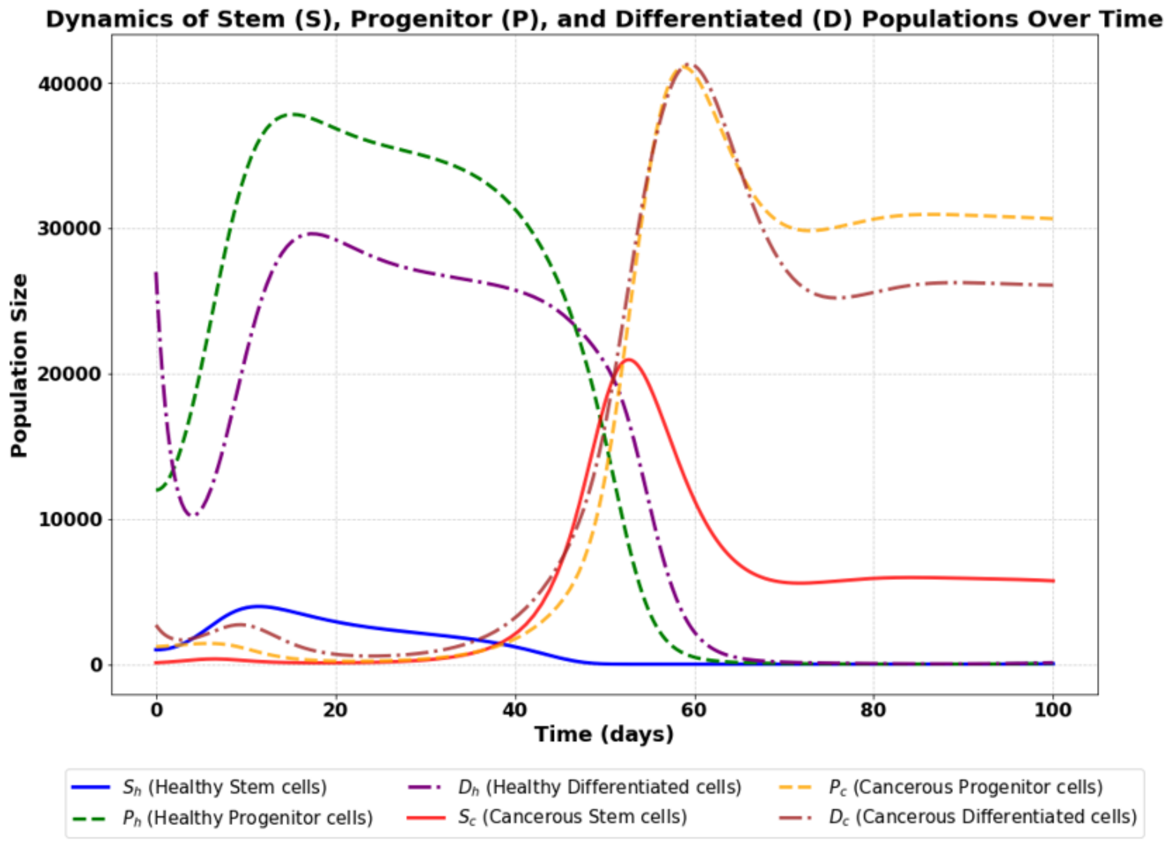

Figure 4: Progression of MDS dynamics between healthy and mutated cell populations under inflammation.

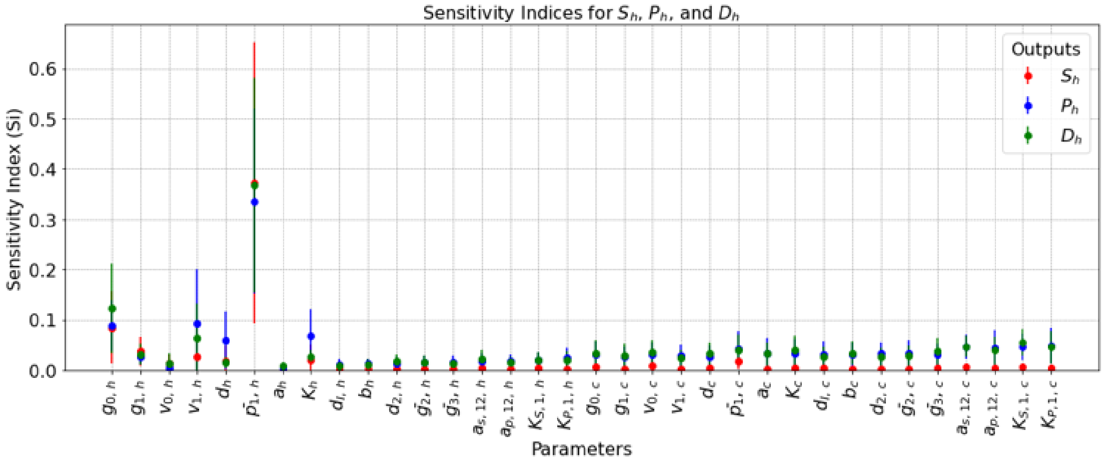

Figure 5: Sensitivity analysis Si values for the healthy S, P and D populations.

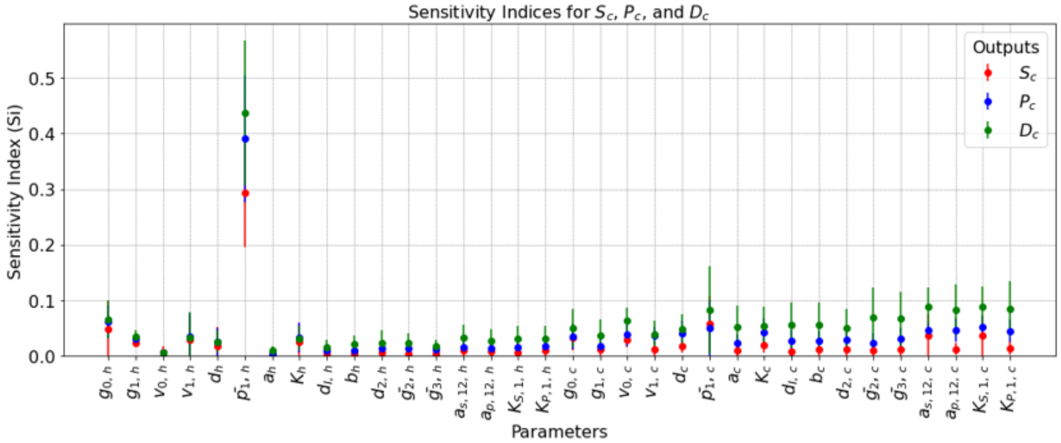

Figure 6: Sensitivity analysis Si values for the cancerous S, P and D populations.

Table 3: Parameter nominal values of the clonal evolution model

| Healthy Population | | Cancerous Population | |
|---|---|---|---|
| **Parameter** | **Value** | **Parameter** | **Value** |
| $v_{0,\,h}$ [d$^{-1}$] | 0.53 | $v_{0,\,c}$ [d$^{-1}$] | 0.53 |
| $v_{1,\,h}$ [d$^{-1}$] | 0.49 | $v_{1,\,c}$ [d$^{-1}$] | 0.49 |
| $g_{0,\,h}$ [cells$^{-1}$] | 3.7·10$^{-5}$ | $g_{0,\,c}$ [cells$^{-1}$] | 3.7·10$^{-5}$ |
| $g_{1,\,h}$ [cells$^{-1}$] | 2.8·10$^{-5}$ | $g_{1,\,c}$ [cells$^{-1}$] | 2.8·10$^{-5}$ |
| $d_h$ [d$^{-1}$] | 0.23 | $d_c[d^{-1}]$ | 0.23 |
| $\overline{p}_{1,\,h}$ [-] | 0.818 | $\overline{p}_{1,\,c}$ [-] | 0.818 |
| $\alpha_h$ [d$^{-1}$] | 0.9 | $\alpha_c$ [d$^{-1}$] | 0.9 |
| $K_h$ [-] | 3 | $K_c$ [-] | 3 |
| $d_{1,\,h}$ [d$^{-1}$] | 20 | $d_{1,\,c}$ [d$^{-1}$] | 20 |
| $\beta_h$ [d$^{-1}$] | 0.8 | $\beta_c$ [d$^{-1}$] | 0.8 |
| $d_{2,\,h}$ [d$^{-1}$] | 0.1 | $d_{2,\,c}$ [d$^{-1}$] | 0.1 |
| $\overline{g}_{2,\,h}$ [cells$^{-1}$] | 0.9·10$^{-5}$ | $\overline{g}_{2,\,c}$ [cells$^{-1}$] | 0.5·10$^{-5}$ |
| $\overline{g}_{3,\,h}$ [cells$^{-1}$] | 0.5·10$^{-8}$ [-] | $\overline{g}_{3,\,c}$ [cells$^{-1}$] | 0.5·10$^{-4}$ |
| $a_{s,\,12,\,h}$ [-] | 0.5 | $a_{s,\,12,\,c}$ [-] | 1.5 |
| $a_{p,\,12,\,h}$ [-] | 0.5 | $a_{p,\,12,\,c}$ [-] | 1.5 |
| $K_{s,\,1,\,h}$ [-] | 5000 | $K_{s,\,1,\,c}$ [-] | 5000 |
| $K_{p,\,1,\,h}$ [-] | 16000 | $K_{p,\,1,\,c}$ [-] | 36000 |
| $S_{0,\,h}$ [cells] | 1000 | $S_{0,\,c}$ [cells] | 100 |
| $P_{0,\,h}$ [cells] | 12000 | $P_{0,\,c}$ [cells] | 1200 |
| $D_{0,\,h}$ [cells] | 27000 | $D_{0,\,c}$ [cells] | 2700 |
